# Scaling pattern of the carnivoran hind limb: Main deviations from a conservative pattern

**DOI:** 10.1101/2022.08.12.503763

**Authors:** Eloy Gálvez-López, Adrià Casinos

**Affiliations:** PalaeoHub, Department of Archaeology, University of York, Wentworth Way, YO10 5DD Heslington, York, United Kingdom; University of Barcelona, Department of Evolutionary Biology, Ecology and Environmental Sciences (BEECA), Barcelona, Spain

**Keywords:** biomechanics, Carnivora, differential scaling, hind limb, locomotor type, phylogenetically independent contrasts, scaling

## Abstract

The scaling pattern of the hind limb in Carnivora was determined using a sample of 13 variables measured on the femur, tibia, and calcaneus, of 429 specimens belonging to 141 species. Standardized major axis regressions on body mass were calculated for all variables, using both traditional regression methods and phylogenetically independent contrasts (PIC). Significant differences were found between the allometric slopes obtained with traditional and PIC regression methods, emphasizing the need to take into account phylogenetic relatedness in scaling studies. Overall, the scaling of the carnivoran hind limb conformed to geometric similarity, although some deviations from its predictions (including differential scaling) were detected, especially in relation with swimming adaptations. The scaling pattern of several phyletic lines and locomotor habits within Carnivora was also determined. Significant deviations from the scaling pattern of the order were found in some phyletic lines, but not in the locomotor habit subsamples. This suggests that the scaling of the carnivoran hind limb is both more heavily influenced by phylogenetic relatedness than by locomotor specializations, and more conservative than that of the forelimb. Finally, together with our previous work on the carnivoran forelimb, the results of the present study suggest that, in large non-aquatic carnivorans, size-related increases in bone stresses are compensated primarily by limb posture changes instead of by modifying limb bone scaling. However, increasing bone robusticity might also occur in the forelimb in response to the heavier stresses acting on the forelimbs due to asymmetrical body weight distribution.

## Introduction

The morphofunctional properties of the bones and muscles of animals change at different rates as their size increases, which is known as scaling (Schmidt-Nielsen, 1984; Alexander, 2002; Biewener, 2003). In order to understand the consequences of scaling, several hypotheses have been proposed which provide theoretical values to these rates of change based on different biomechanical constraints. For instance, the geometric similarity hypothesis derives from the notion that, if all linear dimensions of an object are multiplied by a constant, its volume increases by the cube of this constant, which translates into linear dimensions being proportional to body mass^0.33^ (e.g. Alexander, 2002). On the other hand, the elastic similarity hypothesis proposes that, for different-sized animals to be able to withstand similarly the effect of gravity, their lengths should be proportional to body mass^0.25^ and their diameters to body mass^0.375^ (McMahon, 1973). However, empirical evidence has shown that neither of these hypotheses adequately describes the scaling pattern of mammalian bones (Bou et al., 1987; Bertram & Biewener, 1990; Christiansen, 1999*a*,*b*; Carrano, 2001; Llorens et al., 2001; Lilje et al., 2003; Casinos et al., 2012; Gálvez-López & Casinos, 2022). This is probably related to factors such as phylogenetic constraints to bone morphology (Flynn et al., 1988; Bertram & Biewener, 1990; Day & Jayne, 2007), adaptations to particular locomotor patterns (Bou et al., 1987), or the different biomechanical requirements of locomotion in large and small mammals (Fischer & Blickhan, 2006), which can cause deviations from the theoretical values proposed by the different similarity hypotheses (or even non-linear relationships between those morphofunctional properties and body mass, which is known as differential scaling or complex allometry; Economos, 1983; Bertram & Biewener, 1990; Silva, 1998; Christiansen, 1999a,b; Carrano, 2001).

In order to clarify the effect of these factors on the scaling of the appendicular skeleton, the present work complements a previous work on the scaling of the carnivoran forelimb (Gálvez-López & Casinos, 2022) by presenting here the results obtained for the carnivoran hind limb, as well as an interlimb comparison. The aims of the present study are thus: 1) to determine the scaling pattern of the carnivoran hind limb, and to assess whether differential scaling can be found in this pattern; 2) to analyze whether the main phyletic lines (families) within Carnivora deviated from it, and if so, then how; and 3) to test whether particular locomotor habits within Carnivora cause deviations from the general scaling pattern for the order.

The order Carnivora was chosen because it is a monophyletic group spanning a size range of four orders of magnitude, and presenting one of the widest locomotor diversities among mammals (Van Valkenburgh, 1987; Bertram & Biewener, 1990; Wilson & Mittermeier, 2009; Nyakatura & Bininda-Emonds, 2012).

## Material and Methods

The sample for the hind limb consisted of 429 specimens from 141 species of Carnivora (Table 1). Table 2 describes the locomotor categories used in this study, which represent the locomotor specialization of each species (i.e., the main locomotor habit of each species). For each specimen, measurements were taken on the femur, tibia, and calcaneus. The variables have already been described in the Supplementary Information of Gálvez-López (2021) but are repeated here in Table 3 for simplicity.

**Table 1.**
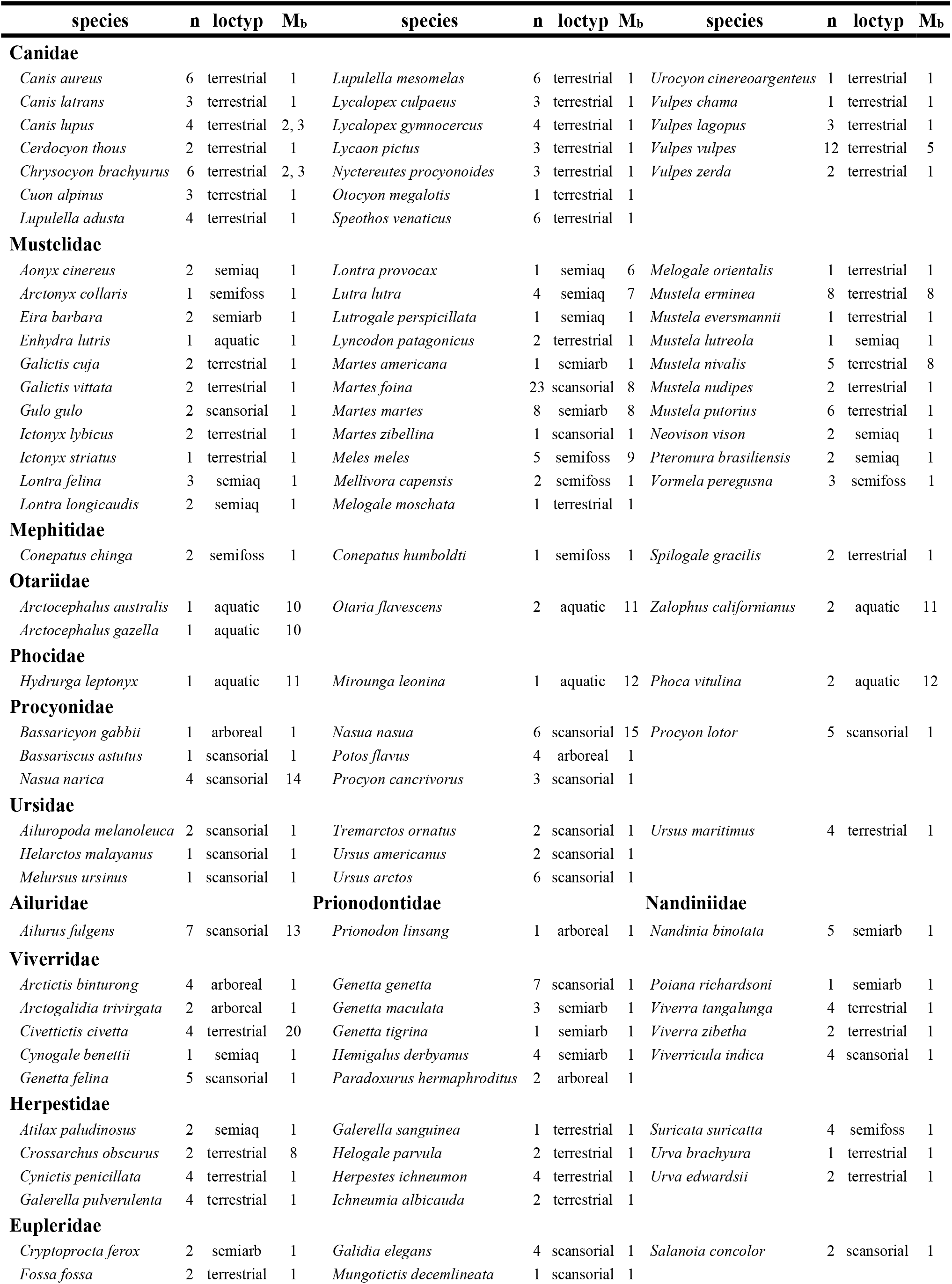
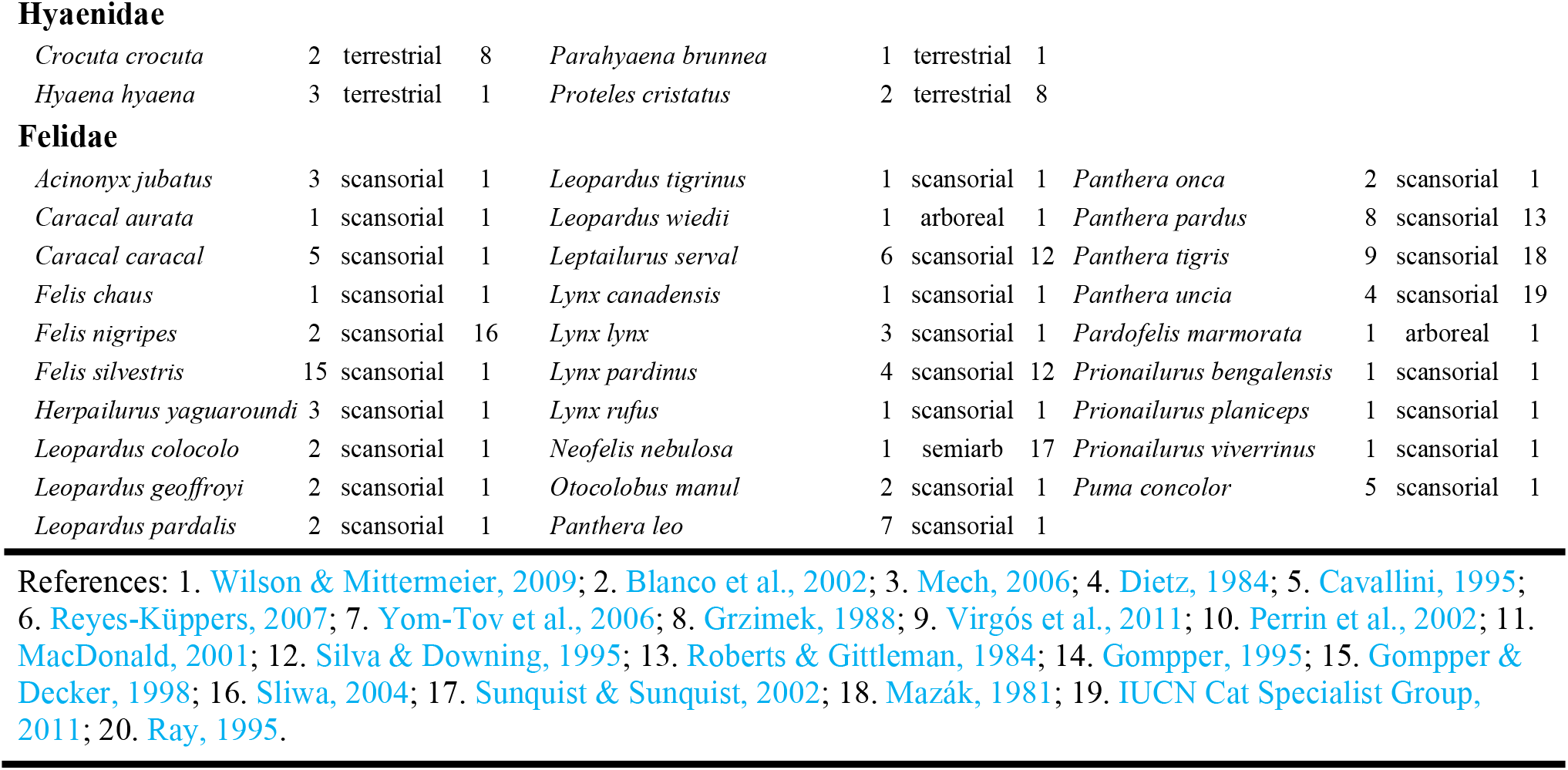
Species measured. For each species, the table shows the number of measured specimens (n), the assigned category for locomotor type (loctyp), and the references from which the mean body mass value for that species was taken (**M_b_**). See Table 2 for a description of locomotor type categories. Abbreviations: semiaq, semiaquatic; semiarb, semiarboreal; semifoss, semifossorial.

| species | n | loctyp | $M_b$ | species | n | loctyp | $M_b$ | species | n | loctyp | $M_b$ |
| --- | --- | --- | --- | --- | --- | --- | --- | --- | --- | --- | --- |
| **Canidae** |  |  |  |  |  |  |  |  |  |  |  |
| *Canis aureus* | 6 | terrestrial | 1 | *Lupulella mesomelas* | 6 | terrestrial | 1 | *Urocyon cinereoargenteus* | 1 | terrestrial | 1 |
| *Canis latrans* | 3 | terrestrial | 1 | *Lycalopex culpaeus* | 3 | terrestrial | 1 | *Vulpes chama* | 1 | terrestrial | 1 |
| *Canis lupus* | 4 | terrestrial | 2, 3 | *Lycalopex gymnocercus* | 4 | terrestrial | 1 | *Vulpes lagopus* | 3 | terrestrial | 1 |
| *Cerdocyon thous* | 2 | terrestrial | 1 | *Lycaon pictus* | 3 | terrestrial | 1 | *Vulpes vulpes* | 12 | terrestrial | 5 |
| *Chrysocyon brachyurus* | 6 | terrestrial | 2, 3 | *Nyctereutes procyonoides* | 3 | terrestrial | 1 | *Vulpes zerda* | 2 | terrestrial | 1 |
| *Cuon alpinus* | 3 | terrestrial | 1 | *Otocyon megalotis* | 1 | terrestrial | 1 |  |  |  |  |
| *Lupulella adusta* | 4 | terrestrial | 1 | *Speothos venaticus* | 6 | terrestrial | 1 |  |  |  |  |
| **Mustelidae** |  |  |  |  |  |  |  |  |  |  |  |
| *Aonyx cinereus* | 2 | semiaq | 1 | *Lontra provocax* | 1 | semiaq | 6 | *Melogale orientalis* | 1 | terrestrial | 1 |
| *Arctonyx collaris* | 1 | semifoss | 1 | *Lutra lutra* | 4 | semiaq | 7 | *Mustela erminea* | 8 | terrestrial | 8 |
| *Eira barbara* | 2 | semiarb | 1 | *Lutrogale perspicillata* | 1 | semiaq | 1 | *Mustela eversmannii* | 1 | terrestrial | 1 |
| *Enhydra lutris* | 1 | aquatic | 1 | *Lyncodon patagonicus* | 2 | terrestrial | 1 | *Mustela lutreola* | 1 | semiaq | 1 |
| *Galictis cuja* | 2 | terrestrial | 1 | *Martes americana* | 1 | semiarb | 1 | *Mustela nivalis* | 5 | terrestrial | 8 |
| *Galictis vittata* | 2 | terrestrial | 1 | *Martes foina* | 23 | scansorial | 8 | *Mustela nudipes* | 2 | terrestrial | 1 |
| *Gulo gulo* | 2 | scansorial | 1 | *Martes martes* | 8 | semiarb | 8 | *Mustela putorius* | 6 | terrestrial | 1 |
| *Ictonyx lybicus* | 2 | terrestrial | 1 | *Martes zibellina* | 1 | scansorial | 1 | *Neovison vison* | 2 | semiaq | 1 |
| *Ictonyx striatus* | 1 | terrestrial | 1 | *Meles meles* | 5 | semifoss | 9 | *Pteronura brasiliensis* | 2 | semiaq | 1 |
| *Lontra felina* | 3 | semiaq | 1 | *Mellivora capensis* | 2 | semifoss | 1 | *Vormela peregusna* | 3 | semifoss | 1 |
| *Lontra longicaudis* | 2 | semiaq | 1 | *Melogale moschata* | 1 | terrestrial | 1 |  |  |  |  |
| **Mephitidae** |  |  |  |  |  |  |  |  |  |  |  |
| *Conepatus chinga* | 2 | semifoss | 1 | *Conepatus humboldti* | 1 | semifoss | 1 | *Spilogale gracilis* | 2 | terrestrial | 1 |
| **Otariidae** |  |  |  |  |  |  |  |  |  |  |  |
| *Arctocephalus australis* | 1 | aquatic | 10 | *Otaria flavescens* | 2 | aquatic | 11 | *Zalophus californianus* | 2 | aquatic | 11 |
| *Arctocephalus gazella* | 1 | aquatic | 10 |  |  |  |  |  |  |  |  |
| **Phocidae** |  |  |  |  |  |  |  |  |  |  |  |
| *Hydrurga leptonyx* | 1 | aquatic | 11 | *Mirounga leonina* | 1 | aquatic | 12 | *Phoca vitulina* | 2 | aquatic | 12 |
| **Procyonidae** |  |  |  |  |  |  |  |  |  |  |  |
| *Bassaricyon gabbii* | 1 | arboreal | 1 | *Nasua nasua* | 6 | scansorial | 15 | *Procyon lotor* | 5 | scansorial | 1 |
| *Bassariscus astutus* | 1 | scansorial | 1 | *Potos flavus* | 4 | arboreal | 1 |  |  |  |  |
| *Nasua narica* | 4 | scansorial | 14 | *Procyon cancrivorus* | 3 | scansorial | 1 |  |  |  |  |
| **Ursidae** |  |  |  |  |  |  |  |  |  |  |  |
| *Ailuropoda melanoleuca* | 2 | scansorial | 1 | *Tremarctos ornatus* | 2 | scansorial | 1 | *Ursus maritimus* | 4 | terrestrial | 1 |
| *Helarctos malayanus* | 1 | scansorial | 1 | *Ursus americanus* | 2 | scansorial | 1 |  |  |  |  |
| *Melursus ursinus* | 1 | scansorial | 1 | *Ursus arctos* | 6 | scansorial | 1 |  |  |  |  |
| **Ailuridae** |  |  |  |  |  |  |  |  |  |  |  |
| *Ailurus fulgens* | 7 | scansorial | 13 | **Prionodontidae** |  |  |  | **Nandiniidae** |  |  |  |
|  |  |  |  | *Prionodon linsang* | 1 | arboreal | 1 | *Nandinia binotata* | 5 | semiarb | 1 |
| **Viverridae** |  |  |  |  |  |  |  |  |  |  |  |
| *Arctictis binturong* | 4 | arboreal | 1 | *Genetta genetta* | 7 | scansorial | 1 | *Poiana richardsoni* | 1 | semiarb | 1 |
| *Arctogalidia trivirgata* | 2 | arboreal | 1 | *Genetta maculata* | 3 | semiarb | 1 | *Viverra tangalunga* | 4 | terrestrial | 1 |
| *Civettictis civetta* | 4 | terrestrial | 20 | *Genetta tigrina* | 1 | semiarb | 1 | *Viverra zibetha* | 2 | terrestrial | 1 |
| *Cynogale benettii* | 1 | semiaq | 1 | *Hemigalus derbyanus* | 4 | semiarb | 1 | *Viverricula indica* | 4 | scansorial | 1 |
| *Genetta felina* | 5 | scansorial | 1 | *Paradoxurus hermaphroditus* | 2 | arboreal | 1 |  |  |  |  |
| **Herpestidae** |  |  |  |  |  |  |  |  |  |  |  |
| *Atilax paludinosus* | 2 | semiaq | 1 | *Galerella sanguinea* | 1 | terrestrial | 1 | *Suricata suricatta* | 4 | semifoss | 1 |
| *Crossarchus obscurus* | 2 | terrestrial | 8 | *Helogale parvula* | 2 | terrestrial | 1 | *Urva brachyura* | 1 | terrestrial | 1 |
| *Cynictis penicillata* | 4 | terrestrial | 1 | *Herpestes ichneumon* | 4 | terrestrial | 1 | *Urva edwardsii* | 2 | terrestrial | 1 |
| *Galerella pulverulenta* | 4 | terrestrial | 1 | *Ichneumia albicauda* | 2 | terrestrial | 1 |  |  |  |  |
| **Eupleridae** |  |  |  |  |  |  |  |  |  |  |  |
| *Cryptoprocta ferox* | 2 | semiarb | 1 | *Galidia elegans* | 4 | scansorial | 1 | *Salanoia concolor* | 2 | scansorial | 1 |
| *Fossa fossa* | 2 | terrestrial | 1 | *Mungotictis decemlineata* | 1 | scansorial | 1 |  |  |  |  |

Hyaenidae
|  |  |  |  |  |  |  |  |
| --- | --- | --- | --- | --- | --- | --- | --- |
| *Crocuta crocuta* | 2 | terrestrial | 8 | *Parahyaena brunnea* | 1 | terrestrial | 1 |
| *Hyaena hyaena* | 3 | terrestrial | 1 | *Proteles cristatus* | 2 | terrestrial | 8 |
| Felidae |  |  |  |  |  |  |  |  |  |  |  |
| *Acinonyx jubatus* | 3 | scansorial | 1 | *Leopardus tigrinus* | 1 | scansorial | 1 | *Panthera onca* | 2 | scansorial | 1 |
| *Caracal aurata* | 1 | scansorial | 1 | *Leopardus wiedii* | 1 | arboreal | 1 | *Panthera pardus* | 8 | scansorial | 13 |
| *Caracal caracal* | 5 | scansorial | 1 | *Leptailurus serval* | 6 | scansorial | 12 | *Panthera tigris* | 9 | scansorial | 18 |
| *Felis chaus* | 1 | scansorial | 1 | *Lynx canadensis* | 1 | scansorial | 1 | *Panthera uncia* | 4 | scansorial | 19 |
| *Felis nigripes* | 2 | scansorial | 16 | *Lynx lynx* | 3 | scansorial | 1 | *Pardofelis marmorata* | 1 | arboreal | 1 |
| *Felis silvestris* | 15 | scansorial | 1 | *Lynx pardinus* | 4 | scansorial | 12 | *Prionailurus bengalensis* | 1 | scansorial | 1 |
| *Herpailurus yaguaroundi* | 3 | scansorial | 1 | *Lynx rufus* | 1 | scansorial | 1 | *Prionailurus planiceps* | 1 | scansorial | 1 |
| *Leopardus colocolo* | 2 | scansorial | 1 | *Neofelis nebulosa* | 1 | semi-arb | 17 | *Prionailurus viverrinus* | 1 | scansorial | 1 |
| *Leopardus geoffroyi* | 2 | scansorial | 1 | *Otocolobus manul* | 2 | scansorial | 1 | *Puma concolor* | 5 | scansorial | 1 |
| *Leopardus pardalis* | 2 | scansorial | 1 | *Panthera leo* | 7 | scansorial | 1 |  |  |  |  |

References: 1. Wilson & Mittermeier, 2009; 2. Blanco et al., 2002; 3. Mech, 2006; 4. Dietz, 1984; 5. Cavallini, 1995; 6. Reyes-Küppers, 2007; 7. Yom-Tov et al., 2006; 8. Grzimek, 1988; 9. Virgós et al., 2011; 10. Perrin et al., 2002; 11. MacDonald, 2001; 12. Silva & Downing, 1995; 13. Roberts & Gittleman, 1984; 14. Gompper, 1995; 15. Gompper & Decker, 1998; 16. Sliwa, 2004; 17. Sunquist & Sunquist, 2002; 18. Mazák, 1981; 19. IUCN Cat Specialist Group, 2011; 20. Ray, 1995.

**Table 2.**
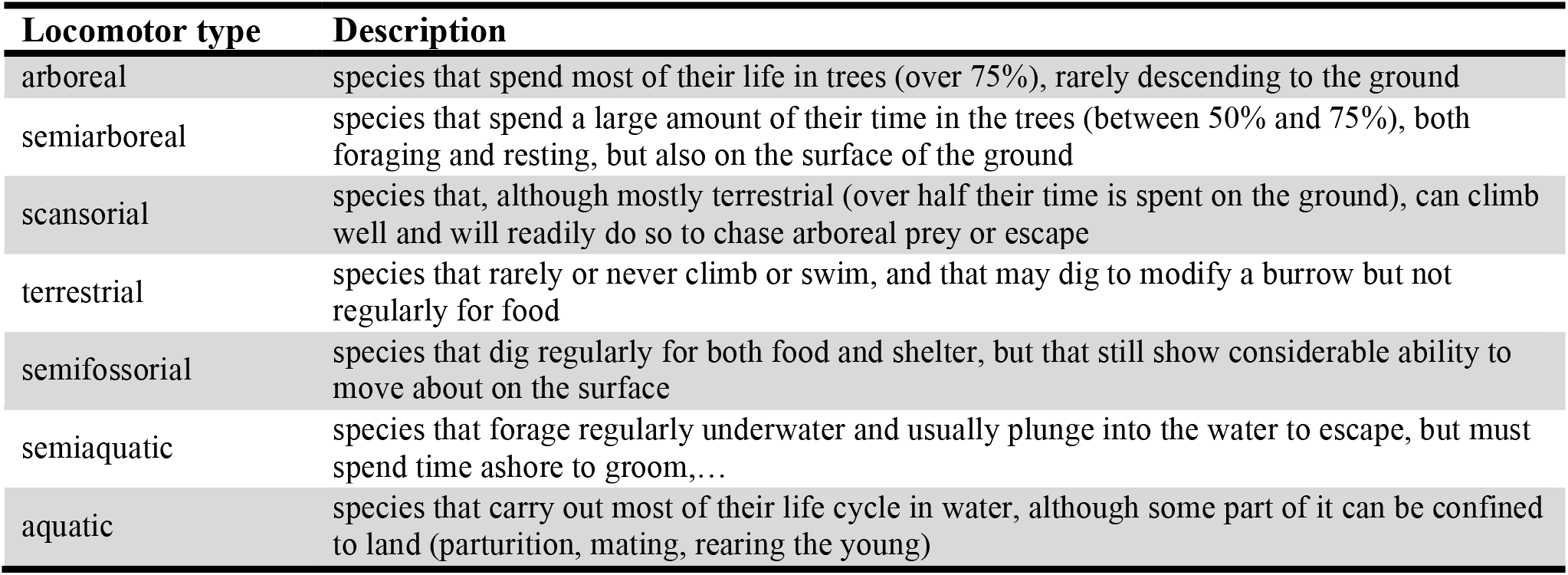
Locomotor type categories. Locomotor type categories were adapted from previous works on the relatioship between locomotor behavior and forelimb morphology (Eisenberg, 1981; Van Valkenburgh, 1985, 1987).

| Locomotor type | Description |
| --- | --- |
| arboreal | species that spend most of their life in trees (over 75%), rarely descending to the ground |
| semi-arboreal | species that spend a large amount of their time in the trees (between 50% and 75%), both foraging and resting, but also on the surface of the ground |
| scansorial | species that, although mostly terrestrial (over half their time is spent on the ground), can climb well and will readily do so to chase arboreal prey or escape |
| terrestrial | species that rarely or never climb or swim, and that may dig to modify a burrow but not regularly for food |
| semifossorial | species that dig regularly for both food and shelter, but that still show considerable ability to move about on the surface |
| semiaquatic | species that forage regularly underwater and usually plunge into the water to escape, but must spend time ashore to groom,... |
| aquatic | species that carry out most of their life cycle in water, although some part of it can be confined to land (parturition, mating, rearing the young) |

**Table 3.**
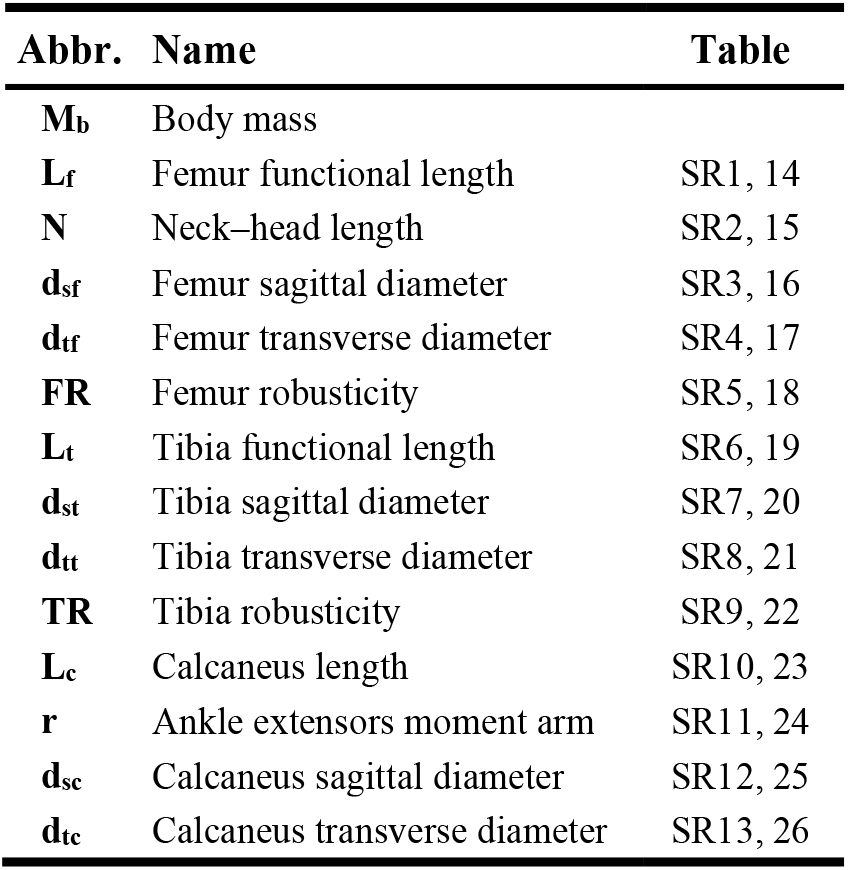
Variable names and their abbreviations. For each variable, it is also indicated which Supplementary Tables show the regression results.

| Abbr. | Name | Table |
| --- | --- | --- |
| **M<sub>b</sub>** | Body mass |  |
| **L<sub>f</sub>** | Femur functional length | SR1, 14 |
| **N** | Neck–head length | SR2, 15 |
| **d<sub>sf</sub>** | Femur sagittal diameter | SR3, 16 |
| **d<sub>tf</sub>** | Femur transverse diameter | SR4, 17 |
| **FR** | Femur robusticity | SR5, 18 |
| **L<sub>t</sub>** | Tibia functional length | SR6, 19 |
| **d<sub>st</sub>** | Tibia sagittal diameter | SR7, 20 |
| **d<sub>tt</sub>** | Tibia transverse diameter | SR8, 21 |
| **TR** | Tibia robusticity | SR9, 22 |
| **L<sub>c</sub>** | Calcaneus length | SR10, 23 |
| **r** | Ankle extensors moment arm | SR11, 24 |
| **d<sub>sc</sub>** | Calcaneus sagittal diameter | SR12, 25 |
| **d<sub>tc</sub>** | Calcaneus transverse diameter | SR13, 26 |

The 13 studied variables included 11 linear measurements and 2 bone robusticities (**FR**, **TR**), and are summarized in Table 3. The linear measurements could be subdivided into bone lengths (represented as ***L_x_***, where x indicates each particular bone, e.g. **L_f_** for femur length), bone diameters (***d_sx_**, **d_tx_***), and other measurements (**N**, **r**). Bone robusticities were calculated dividing sagittal diameter by bone length (***XR*** = **d_sx_** / **L_x_**). Since the scapula has been shown to be the main propulsive element of the forelimb (Fischer et al., 2002; Lilje & Fischer, 2001), it is thus considered here the most proximal segment of the forelimb and, correspondingly, the functional homologue of the femur.

Following Gálvez-López & Casinos (2022), all variables were regressed to body mass (**M_b_**) using the standardised major axis method (SMA) and assuming the power equation (*y* = *a · x^b^*, Eq. 1) for all of them. All regressions were calculated using PAST (Hammer et al. 2001), and 95% confidence intervals were obtained for both the coefficient (*a*) and the allometric exponent (*b_trad_*). Furthermore, all the SMA regression slopes using phylogenetically independent contrasts (PIC) were also calculated, since the hierarchical sequence of interspecific data introduces a phylogenetic signal that could cause correlation of the error terms due to the lack of independence among species (Felsenstein, 1985; Grafen, 1989; Harvey & Pagel, 1991; Christiansen, 2002a, *b*). PIC regression slopes were calculated using the PDAP: PDTREE module of Mesquite (Maddison & Maddison, 2010; Midford et al., 2010). The structure of the phylogenetic tree used in the present study is discussed elsewhere (Gálvez-López & Casinos, 2022). The PIC slopes (*b_PIC_*) were compared to *b_trad_* values with an F-test (p < 0.05) to assess whether the phylogenetic signal had any effect on the results.

As in the previous study (Gálvez-López & Casinos, 2022), for each variable and methodology (traditional and PIC), separate regressions were calculated for the whole sample, the fissiped subsample, and also for several subsamples by family and locomotor type. Regressions were not calculated for any subsample with a sample size lower than 5, which was the case for Hyaenidae, Mephitidae, Phocidae, Otariidae, Prionodontidae, the monotypic families (Ailuridae, Nandiniidae), and Eupleridae when using PIC regression, plus a few other groups in the case of calcaneal variables (Ursidae, Herpestidae, Eupleridae, and aquatic and semifossorial carnivorans, plus semiarboreal carnivorans when using PIC regression). Then, all allometric exponents were compared to the theoretical values proposed by the geometric similarity hypothesis (all variables **∝ M_b_^0.33^**, except for **FR**, **TR** ∝ **M_b_**^0^) and the elastic similarity hypothesis (***L_x_*, N, r ∝ M_b_**^0.25^; ***d_sx_, d_tx_* ∝ M_b_**^0.375^; **FR, TR ∝ M_b_**^0.125^). Allometric exponents were considered to deviate significantly from the predictions of any similarity hypothesis when their 95%CI did not include the corresponding theoretical value. Furthermore, for each variable, allometric exponents were compared between the whole sample and the fissiped subsample, and between the different family and locomotor type subsamples.

Finally, also for each variable and each subsample, the presence of differential scaling was assessed using the model proposed by Jolicoeur (1989; ln *y* = ln *A – C* · (ln ***x***_*max*_ – ln ***x***)^*D*^, Eq. 2), which was fitted with SPSS for Windows (release 15.0.1 2006; SPSS Inc., Chicago, IL, USA), and 95% confidence intervals were calculated for all parameters.

## Results

Supplementary Tables SR1 through SR13 show the regression results for each variable. As observed in previous studies comparing traditional and PIC regressions (Christiansen, 2002a, b; Gálvez-López & Casinos, 2012, 2022), the correlation coefficients (R) from the PIC analyses were lower than those from traditional regressions in most cases, which sometimes resulted in regressions no longer being significant (e.g. Table SR13). Some authors have attributed this phenomenon to a higher risk of type I errors (i.e., indicating a significant correlation between two variables when there was none) when the effect of phylogeny is neglected in correlation analyses (Grafen, 1989; Christiansen, 2002a). In several cases, however, *R* actually increased after taking into account the effect of phylogeny, which in some cases resulted in regressions becoming significant (e.g. Table SR1). Branch lengths ought to be transformed in most cases before performing the PIC regressions (Table S1).

### Whole sample vs. Fissiped subsample

As in the forelimb (Gálvez-López & Casinos, 2022), removal of Pinnipedia from the sample caused a generalized increase of the allometric exponents (especially when using traditional regression methods), although this increase was only significant for **L_f_**, **N**, **d_sf_**, and **d_st_**, and then only for *btrad* (Tables SR1–SR3, SR7). Overall, PIC slopes tended to be lower than those obtained using traditional regression methods, being significantly lower for **L_f_** and **L_t_** in the fissiped subsample (Tables SR1, SR6). Significant differences between the allometric exponents obtained using traditional and PIC regressions were also found for the carnivoran forelimb (Gálvez-López & Casinos, 2022), but not in other previous studies comparing both methodologies (Christiansen, 2002b; Christiansen & Adolfssen, 2005; Gálvez-López & Casinos, 2012).

Regarding conformity with the similarity hypotheses, Table 4 summarizes the percentage of linear measurements (i.e., no ratios: **FR**, **TR**) that conform to each similarity hypothesis in both the whole sample and the fissiped subsample, and also using either traditional regression methods or PIC. Contrary to the results obtained for the forelimb (Gálvez-López & Casinos, 2022), the scaling pattern of the hind limb in Carnivora conformed clearly to the geometric similarity hypothesis. However, the removal of Pinnipedia from the sample produced conflicting results between both methodologies (Table 4): while traditional regression results indicated low conformity to either similarity hypotheses, PIC slopes conformed to the geometric similarity hypothesis, like in the whole sample. Finally, both bone robusticities scaled elastically, whatever the sample or methodology (Tables SR5, SR9).

**Table 4.**
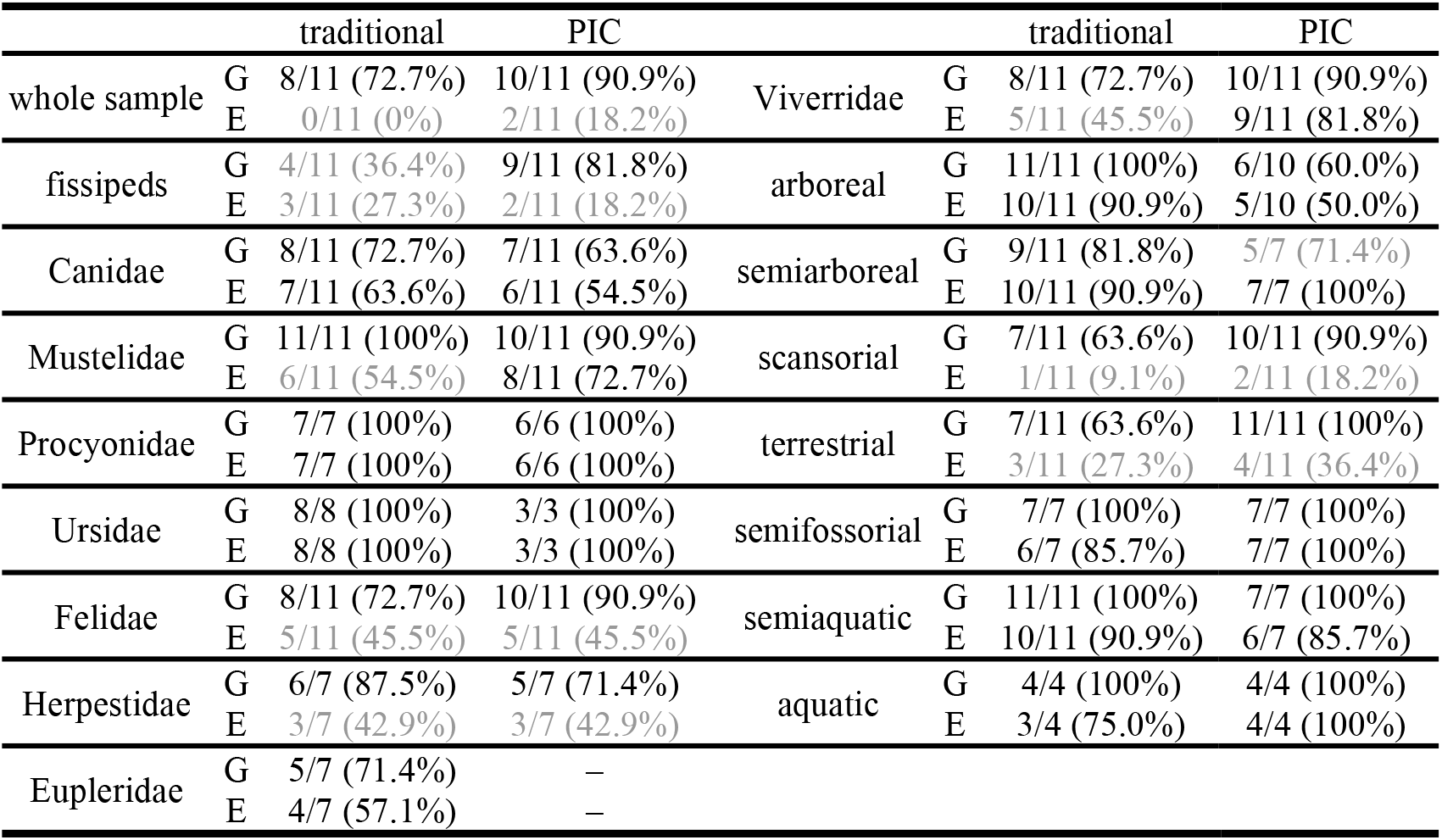
Conformity to the similarity hypotheses summary. For each subsample, the number of linear measurements conforming to geometric (G) or elastic similarity (E) is given, as is the percentage of the significant regressions for that subsample that they represent. Values in grey indicate that the number of variables conforming to a particular similarity hypothesis is either less than half the number of variables, or over 20% lower than the number of variables conforming to the other similarity hypothesis.

### Family subsamples

No significant differences were found between the allometric exponents obtained with each method (Tables SR1–SR13), which agrees with previous studies comparing traditional and PIC regression methods (Christiansen, 2002b; Christiansen & Adolfssen, 2005; Gálvez-López & Casinos, 2012) and with a previous work on forelimb scaling in the same carnivoran families (Gálvez-López & Casinos, 2022).

Like in the whole sample, the scaling pattern of the carnivoran hind limb conformed to geometric similarity in most families (Mustelidae, Felidae, Herpestidae, and Viverridae), although for Mustelidae and Viverridae the elastic similarity hypothesis was also a likely explanation after taking into account the effect of phylogenetic relatedness (Table 4; PIC results). In Ursidae and Procyonidae, however, small sample sizes resulted in low correlation coefficients (R) and 95%CIb wide enough to include the theoretic value for both hypotheses in most of the variables, and thus no similarity hypothesis could be ruled out (Table 4). Finally, in the case of Canidae, conformity to both similarity hypotheses was low, especially considering PIC results.

Regarding bone robusticities, when significant, **TR** always scaled elastically (Table SR9). In the case of **FR**, the slopes calculated using traditional regression methods presented intermediate values between the theoretical values proposed by both similarity hypotheses. PIC slopes for **FR**, however, conformed to the elastic similarity hypothesis (Table SR5). Figure 1 shows comparisons of the allometric exponents between different families for each variable, which are summarized in Table 5. No significant differences between families were found for **TR**. Overall, Canidae and Eupleridae scaled faster that all other families, while Herpestidae and Viverridae present lower allometric slopes than most families.

**Figure 1.**
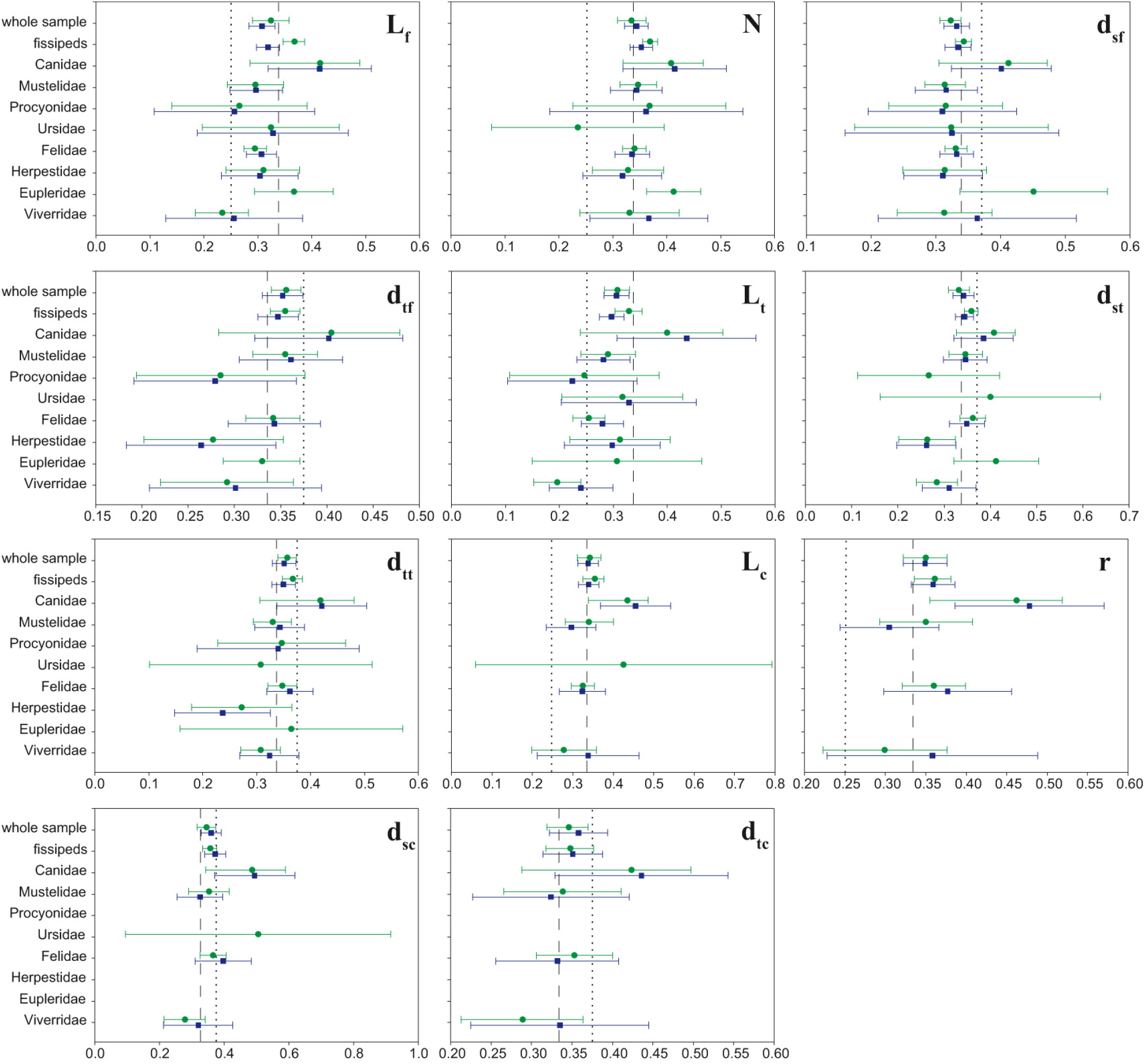
Allometric exponents by family. For each subsample, the allometric exponents obtained using traditional regression methods (green) and phylogenetically independent contrasts (blue), as well as their 95% confidence intervals, are shown. Only the results of significant regressions are presented. The allometric slopes obtained for the whole sample and the fissiped subsample are included as a reference. The dashed line represents the theoretical value proposed by the geometric similarity hypothesis, while the dotted line corresponds to that proposed by the elastic similarity hypothesis. Variable names are listed in Table 3.

**Table 5.**
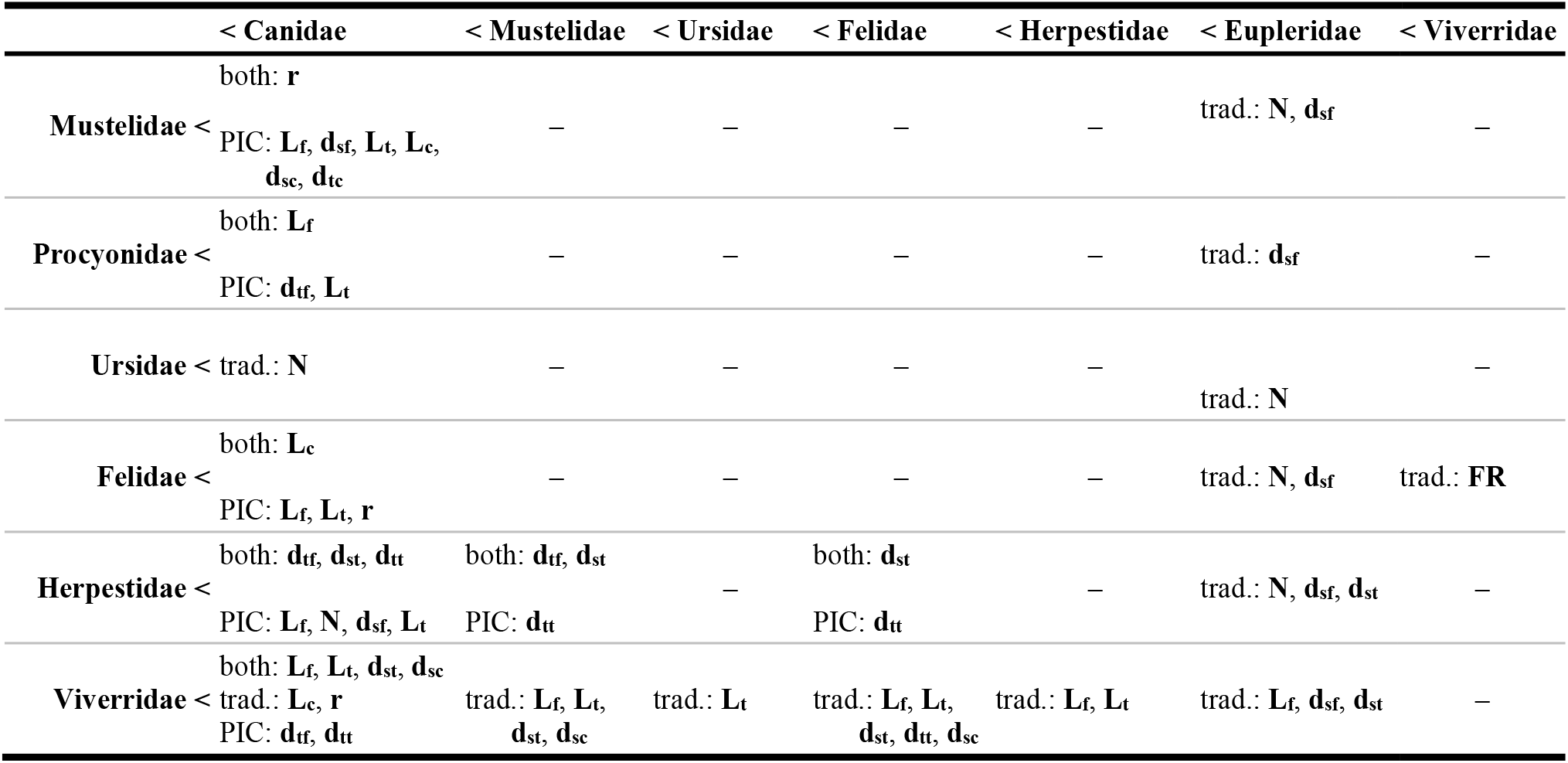
Differences in allometric exponents between families. Rows list families with an allometric exponent (b) significantly lower than the families listed in columns. That can happen when comparing allometric exponents from traditional regression (trad.), phylogenetically independent contrasts (PIC), or when using both methodologies (both). Variable names are listed in Table 3.

### Locomotor type subsamples

Contrary to previous studies comparing traditional and PIC regression methods (Christiansen, 2002b; Christiansen & Adolfssen, 2005; Gálvez-López & Casinos, 2012) but in agreement with a previous work on forelimb scaling in Carnivora (Gálvez-López & Casinos, 2022), significant differences between the allometric exponents obtained with each methodology were observed: for both **L_f_** and **L_t_** traditional slopes were significantly higher than PIC slopes in terrestrial carnivorans (Tables SR1, SR6).

The scaling pattern of scansorial and terrestrial carnivorans conformed better to the geometric similarity hypothesis (Table 4). In the rest of locomotor type subsamples, the 95% CIb were wide enough to include the theoretic value for both hypotheses in most of the variables and thus no similarity hypothesis could be ruled out. However, note that, according to PIC slopes, semiarboreal carnivorans scaled elastically, and that conformity to both similarity hypotheses was low in arboreal carnivorans (due to the particular scaling of their calcaneal variables) (Table 4).

Regarding bone robusticities, when significant, they scaled elastically, except for **FR** in terrestrial carnivorans, which decreased with increasing body mass values (Tables SR5, SR9). Figure 2 shows comparisons of the allometric exponents between different locomotor types for each variable, which are summarized in Table 6. Most significant differences between locomotor types were found either among traditional or PIC slopes, but not both.

**Figure 2.**
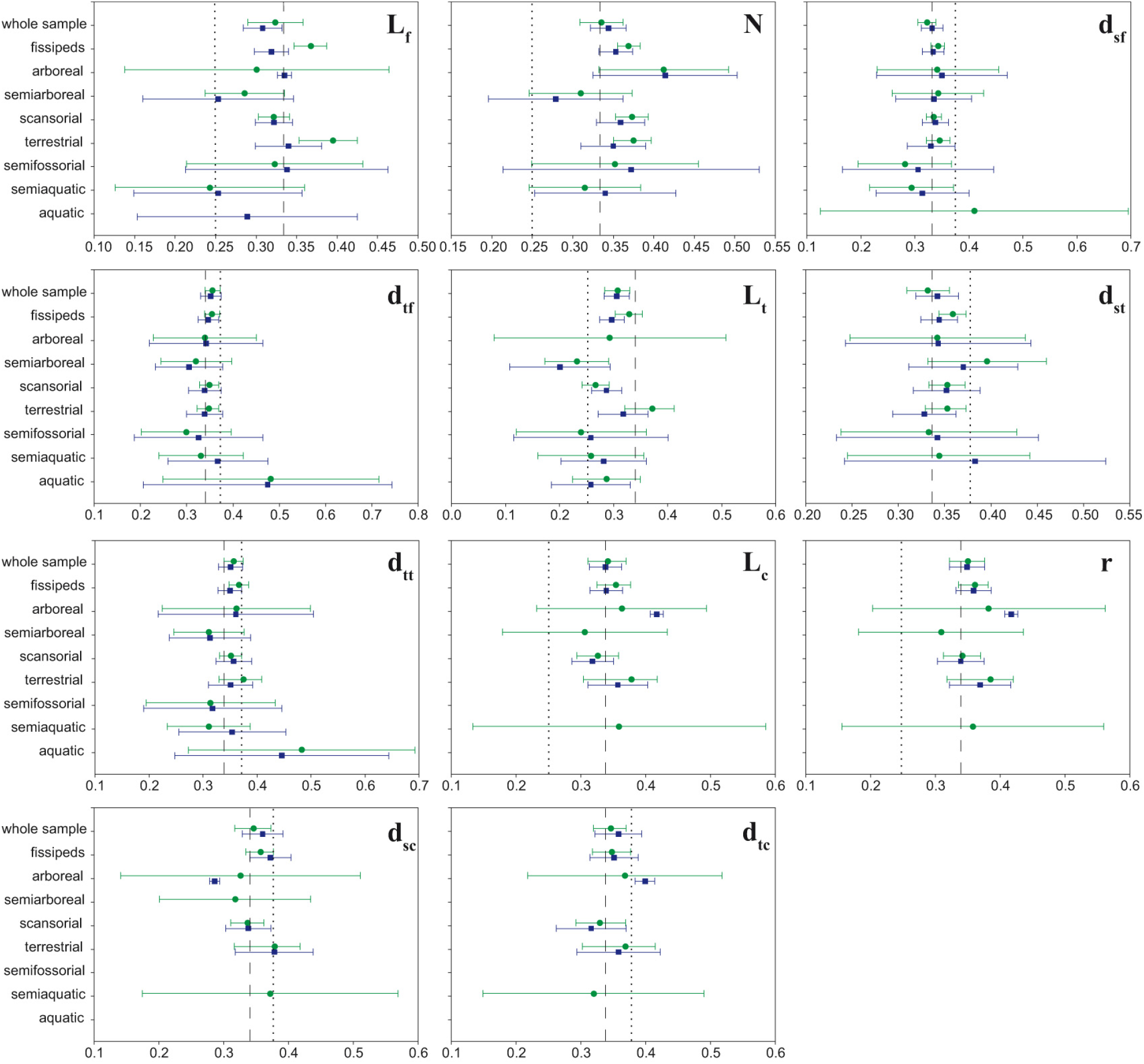
Allometric exponents by locomotor type. For each subsample, the allometric exponents obtained using traditional regression methods (green) and phylogenetically independent contrasts (blue), as well as their 95% confidence intervals, are shown. Only the results of significant regressions are presented. The allometric slopes obtained for the whole sample and the fissiped subsample are included as a reference. The dashed line represents the theoretical value proposed by the geometric similarity hypothesis, while the dotted line corresponds to that proposed by the elastic similarity hypothesis. See Table 2 for a description of locomotor type categories. Variable names are listed in Table 3.

**Table 6.** Differences in allometric exponents between locomotor types. Rows list categories with an allometric exponent (*b*) significantly lower than the categories listed in columns. That can happen when comparing allometric exponents from traditional regression (trad.), phylogenetically independent contrasts (PIC), or when using both methodologies (both). Variable names are listed in Table 3.

|  | < arboreal | < scansorial | < terrestrial | < semiaquatic |
| --- | --- | --- | --- | --- |
| arboreal < | – | PIC: *d<sub>sc</sub>* | PIC: *d<sub>sc</sub>* | – |
| semiarboreal < | both: *N* | – | both: *L<sub>t</sub>*<br>trad.: *L<sub>f</sub>*, *N* | – |
| scansorial < | PIC: *L<sub>c</sub>*, *r*, *d<sub>tc</sub>* | – | trad.: *L<sub>f</sub>*, *L<sub>t</sub>* | – |
| terrestrial < | PIC: *L<sub>c</sub>*, *r* | – | – | trad.: *FR* |
| semifossorial < | – | – | trad.: *L<sub>t</sub>* | – |
| semiaquatic < | trad.: *N* | – | trad.: *L<sub>f</sub>*, *L<sub>t</sub>* | – |
| aquatic < | – | – | trad.: *L<sub>f</sub>*, *L<sub>t</sub>* | – |

### Complex allometry

Results for the test for complex allometry are shown in Supplementary Tables SR14 through SR26.

Evidence for complex allometry was found in half of the variables for the whole sample. In the case of **L_f_**, **N**, **L_t_**, **d_st_**, **L_c_**, and **r**, *D* was significantly higher than 1, indicating that these variables scale faster in small species; while in **d_tf_** *D* was significantly lower than 1, suggesting faster scaling in large species.

As observed for the carnivoran forelimb (Gálvez-López & Casinos, 2022), after removing Pinnipedia from the sample (i.e., fissiped subsample), evidence for complex allometry was rarely recovered. Only for **L_t_** and **L_c_** was *D* still significantly different from 1 (D > 1 in both cases). Furthermore, significant evidence for complex allometry was also found for **TR**, which presented *D* < 1. However, the 95% CI_*D*_ for **TR** included 0, indicating independence from body mass (Gálvez-López & Casinos, 2022).

Overall, significant evidence for complex allometry was scarce in the family subsamples. Differential scaling was found in Mustelidae (**d_sf_**), Herpestidae (**L_f_**, **N**, **L_t_**, **d_tt_**), and Viverridae (**L_f_**, **N**, **d_sf_**, **d_tf_**, **L_t_**, **d_st_**, **d_tc_**), with *D*<1 in all cases. However, as observed for the whole sample when *D*<1, in some cases the 95%CI_*D*_ also included 0, indicating independence from body mass: **N** in Viverridae, and **L_t_** and **d_tt_** in Herpestidae.

In the locomotor type subsamples, significant evidence for complex allometry was even less frequent than in the family subsamples. In all locomotor types but scansorial, when complex allometry was detected, it suggested that large carnivorans scaled faster than small species (i.e., *D* < 1). This was the case for **N** and **d_tf_** in arboreal carnivorans, for **L_f_** in semiarboreal species, and for **TR** in semifossorial carnivorans. In the case of scansorial carnivorans, on the other hand, large species scaled lower (**L_f_**, **d_sf_**, **L_t_**, **L_c_**). As previously observed, in some cases when *D*<1 the 95% CI_*D*_ also included 0. This was the case for **d_tf_** in arboreal carnivorans, and **TR** in semifossorial species.

## Discussion

### Considerations on the scaling pattern of the carnivoran hind limb

Previous studies on hind limb scaling have found scarce conformity to either the geometric or the elastic similarity hypotheses (Bou et al., 1987; Bertram & Biewener, 1990; Cubo & Casinos, 1998; Heinrich & Biknevicius, 1998; Christiansen, 1999a,b; Carrano, 2001; Llorens et al., 2001; Lilje et al., 2003; Casinos et al., 2012). In the present study, however, high conformity to geometric scaling was the norm in most subsamples, especially when considering the results of phylogenetically independent contrasts (PIC; Table 4). Regardless of conformity, significant evidence for complex allometry was found in most of the studied variables, which agrees with previous studies (Economos, 1983; Bertram & Biewener, 1990; Silva, 1998; Christiansen, 1999a,b; Carrano, 2001). Finally, in agreement with previous results on the carnivoran forelimb (Gálvez-López & Casinos, 2022), but contrary to previous studies comparing traditional and PIC regression methods (Christiansen, 2002b; Christiansen & Adolfssen, 2005; Gálvez-López & Casinos, 2012), significant differences between the slopes of both methodologies were found, especially in the fissiped subsample. Thus, in order to avoid any possible artefacts caused by the phylogenetic relatedness of the species in our sample, only the PIC results will be further discussed.

It has been proposed that differential scaling is a consequence of the need of large mammals (over 200kg) to develop progressively more robust limb bones in order to endure the higher bone stresses caused by increasing body mass (Biewener, 1990; Christiansen, 1999a,b; Carrano, 2001). At body mass values under that threshold, postural modifications alone (limb straightening) should reduce the magnitude of those stresses, and thus no changes in bone scaling would be required (Biewener, 2003; Carrano, 2001). Since only a handful of non-aquatic carnivoran species attain such large body sizes, limb straightening should be the primary strategy to reduce bone stresses for most carnivorans, not changing limb bone scaling (i.e., differential scaling). In the present study, however, significant evidence for differential scaling was found in several variables, indicating that hind limb scaling does change with size in Carnivora. As observed previously for the carnivoran forelimb (Gálvez-López & Casinos, 2022), differential scaling was not recovered for most variables after removing Pinnipedia from the sample (i.e., in the fissiped subsample), probably indicating that scaling changes were related to the locomotor specialization of that group (swimming). Furthermore, no significant evidence for differential scaling was found in any of the “large” carnivoran families (Canidae, Felidae, Ursidae; Tables SR14–SR26). Thus, together with our previous work on the carnivoran forelimb (Gálvez-López & Casinos, 2022), the results of the present study suggest that, in large non-aquatic carnivorans, size-related increases in bone stresses are compensated by limb posture changes instead of by modifying limb bone scaling.

As observed in previous studies, femur generally presented higher correlation coefficients than tibia, which has been interpreted as proximal proximal limb segments being more conservative in lengthening with increasing body mass than distal ones (McMahon, 1975; Lilje et al., 2003; Schmidt & Fischer, 2009; Gálvez-López & Casinos, 2022). Interestingly, all subsamples including species adapted to swimming (whole sample; semiaquatic and aquatic carnivorans; freshwater and marine species) deviated from this tendency, presenting more variability in the length of the femur than in the tibia. This is probably related to the limb bone shortening described previously in carnivorans adapted to swimming, which was particularly evident for the femur (Gálvez-López, 2021).

Previous studies had reported scaling differences between femur and tibia (Casinos et al., 1986; Wayne, 1986; Bertram & Biewener, 1990; Raich & Casinos, 1991; Heinrich & Biknevicius, 1998; Christiansen, 1999a; Llorens et al., 2001; Lilje et al., 2003; Casinos et al., 2012). In those studies, the length of the tibia tended to scale slower than the length of the femur. In the present study, although the tibia presented the lowest allometric exponent in most subsamples, significant differences were only found for semiarboreal carnivorans. Furthermore, both femur and tibia length scaled slower than calcaneus length in most subsamples. The scaling of bone diameters also tends to be faster in the femur than in the tibia (Llorens et al., 2001; Lilje et al., 2003; but see Cubo & Casinos, 1998). The results of the present study, however, suggest that both sagittal and transverse diameters of femur and tibia scale no differently from each other in Carnivora. In fact, only for Herpestidae were some differences between bone diameters significant (**dsf** scaled faster than **dtt**; Tables SR12, SR13). Thus, together with our previous study on forelimb scaling (Gálvez-López & Casinos, 2022), the results of the present study suggest that the scaling pattern of the carnivoran hind limb is more conservative than that of the forelimb.

### Phylogeny, locomotor habit and the scaling of the carnivoran hind limb

Overall, the scaling patterns found for the hind limb in the different carnivoran families were similar to the pattern found for the whole order. Only Canidae and Herpestidae deviated significantly from it (Fig. 1). In the case of Canidae, bone lengths (**L_f_**, **L_t_**, **L_c_**) and the moment arm of the ankle extensors (**r**) scaled faster than in the rest of Carnivora; while in Herpestidae several bone diameters presented lower slopes than those of the whole order and the fissiped subsample (**d_tf_**, **d_st_**, **d_tt_**). Furthermore, when comparing the allometric exponents obtained for each variable between families, Canidae and Eupleridae tended to scale faster than other families, while Herpestidae and Viverridae presented lower allometric slopes than most families. In canids this could be explained by size selection, which seems to be one of the main forces driving canid evolution (Wayne, 1986; Gálvez-López & Casinos, 2022). The slow scaling of bone lengths in Viverridae could be related to arboreality, since presenting short limbs has been described in this family as a strategy to increase stability in arboreal supports (Cartmill, 1985). In the case of Herpestidae, the low allometric slopes found for bone diameters could reflect a reduction in fossorial habits with increasing size, since fossorial species tend to have more robust limb bones than less specialized species (Lehmann, 1963; Casinos et al., 1993; Elissamburu & Vizcaíno, 2004; Gálvez-López, 20221). Finally, it should be noted that the wide confidence intervals (95%CIb) obtained for some families could be obscuring further significant deviations from the ordinal scaling pattern (e.g. Procyonidae, Eupleridae).

The lack of significant differences between traditional and PIC slopes in the family subsamples agrees with both a previous study stating that most morphological variability of the appendicular skeleton in Carnivora occurs at the family level (Gálvez-López, 2021), and the results found for the carnivoran forelimb (Gálvez-López & Casinos, 2022). Also in agreement with previous results on the carnivoran forelimb, significant differences between both methodologies were found for several locomotor type subsamples (Gálvez-López & Casinos, 2022).

Previous studies on the scaling of the appendicular skeleton in Carnivora have been restricted to fissiped carnivorans, and thus comparisons with the literature will only be discussed for that subsample. Overall, our results using traditional regression methods tend to agree with those of Bertram & Biewener (1990) regarding conformity to the similarity hypotheses. However, since they regressed bone lengths onto bone diameters, no direct comparison could be made. Similarly, the PIC slopes obtained for both femur and tibia length matched those obtained by Christiansen (1999a). Conformity to the geometric similarity hypothesis was high in Felidae, in agreement with previous studies on the scaling of their appendicular skeleton (Day & Jayne, 2007; Gálvez-López & Casinos, 2012, 2022). Finally, Heinrich & Biknevicius (1998) obtained lower allometric slopes than those proposed by any similarity hypothesis for the hind limbs of scansorial mustelids (i.e., Martinae), which were not recovered in the present study for either Mustelidae or scansorial carnivorans (no specific regressions were carried out for any subfamily).

Contrary to the forelimbs, which are involved in both locomotion and prey capture/handling in Carnivora (Iwaniuk et al., 1999), the hind limbs are merely locomotor. Furthermore, even within their locomotor function, the forelimbs generally perform a wider variety of tasks than the hindlimbs (e.g. semifossorial carnivorans dig exclusively with their forelimbs; Wilson & Mittermeier, 2009). Thus, selective pressures acting on the hind limb are probably similar for all carnivoran species, regardless of locomotor habit, which would explain why locomotor type subsamples did not deviate significantly from the scaling pattern of the whole sample (Figs. 2, 3). Another possible explanation could be that bone scaling was more heavily influenced by phylogenetic relatedness than by other factors (Lilje et al., 2003). This argument is supported by the lower amount of significant differences in the allometric slopes among locomotor type or preferred habitat subsamples than among carnivoran families, especially when considering PIC results (Tables 5, 6), which was also observed in the carnivoran forelimb (Gálvez-López & Casinos, 2022). Finally, it is worth noting that calcaneal variables did deviate from the scaling pattern of the whole sample in arboreal carnivorans. However, judging by the unexpectedly high correlation coefficients and narrow 95%CI_b_’s for such low sample sizes (Tables SR10–SR13), these results are probably spurious and were thus not taken into account.

Similarly to previous results on the carnivoran forelimb (Gálvez-López & Casinos, 2022), significant evidences for complex allometry were found in several variables measured on the carnivoran hind limb. Again, the causes of this differential scaling are hard to ascertain, since it was detected in subsamples with wide and narrow body mass ranges (e.g. the whole sample and Herpestidae), in those including a wide variety of locomotor types (e.g. Viverridae), and in some locomotor type subsamples (e.g. semiarboreal and scansorial carnivorans). Thus, neither body mass range (Economos, 1983; Bertram & Biewener, 1990), varying locomotor requirements (Biewener, 1990, 2003; Christiansen, 1999a,b; Carrano, 2001), nor the inclusion of different locomotor types in the same sample (Gálvez-López & Casinos, 2012), provide a sound explanation for differential scaling in Carnivora.

### Interlimb scaling in Carnivora

Previous interlimb comparisons of the scaling of bone lengths have revealed that the distal forelimb segments (i.e., radius/ulna, metacarpals) scale faster than both the femur and the tibia (Wayne, 1986; Raich & Casinos, 1991; Heinrich & Biknevicius, 1998; Christiansen, 1999a; Llorens et al., 2001; Lilje et al., 2003). However, conflicting results exist on the relationships between the proximal limb segments (i.e., scapula and humerus in the forelimb and femur and tibia in the hind limb). In their study on the scaling of the appendicular skeleton in some scansorial mustelids, Heinrich & Biknevicius (1998) obtained higher allometric slopes for humerus than for femur. Similar results were obtained by Llorens et al. (2001) in Platyrrhina. As Heinrich & Biknevicius (1998) pointed out, this would indicate a greater straightening of the hind limbs with increasing body mass, since pivot height, and thus functional length, is the same for both the forelimbs and the hind limbs in mammals (Fischer & Blickhan, 2006). On the other hand, whereas in Carnivora the length of femur, humerus and ulna scaled no differently from each other, and all of them faster than tibia length (Raich & Casinos, 1991), in Canidae the length of the femur scaled faster than that of both the humerus and the tibia, but no differently from from scapular length (Wayne, 1986). Furthermore, Christiansen (1999a) found no significant differences in the scaling of the length of humerus, femur and tibia in mammals. However, in the same study he described a lower slope for the tibia than for humerus and femur in Carnivora, which was also later observed by Lilje et al. (2003) in Ruminantia. In the present study, both tibia and femur length scale slower (often significantly) than the length of all forelimb bones except for humerus, which suggests that indeed limb straightening is greater in the hind limbs than in the forelimbs. In the case of Ursidae, femur and tibia length scale faster than the length of any forelimb bone but the third metacarpal, which might be interpreted as the forelimbs straighten further than the hind limbs. However, bears have plantigrade feet, and thus the higher slopes for femur and tibia length probably compensate the loss of the distal (metatarsal) segment. The rest of subsamples generally follow the scaling pattern found for Carnivora, although with slight deviations which probably do not affect the limb straightening relationship (e.g. both ulna and radius length scale slower than femur length in semiarboreal mammals).

In the case of bone diameters, previous interlimb comparisons agree on hind limb diameters scaling slower than forelimb diameters (Cubo & Casinos, 1998; Heinrich & Biknevicius, 1998; Llorens et al., 2001; Lilje et al., 2003), which was also recovered for most subsamples in the present study. These results agree with those of Carrano (2001), who stated that hind limb bones are relatively slenderer than forelimb bones. The only exceptions to this trend were 1) the sagittal diameter of the third metacarpal, which scaled significantly slower than femur sagittal diameter in Ursidae, and than tibia transverse diameter in forest-dwelling carnivorans; and 2) radius sagittal diameter, which scaled significantly slower than all hind limb diameters in semiarboreal carnivorans.

Thus, in Carnivora, hind limbs are straightened more with increasing size than forelimbs, while bone robusticity increases faster in the forelimbs than the hind limbs, which would suggest that each pair of limbs presents a different strategy to cope with the higher bone stresses caused by increasing body mass (Biewener, 1990; Christiansen, 1999a,b; Carrano, 2001). In turn, this might be related to the asymmetrical distribution of body weight in most mammals, which imposes heavier loads on the forelimb than on the hind limbs (Schmitt & Lemelin, 2002).

## Acknowledgements

We would like to thank the curators of the Phylogenetisches Museum (Jena), the Museum für Naturkunde (Berlin), the Museu de Ciències Naturals de la Ciutadella (Barcelona), the Muséum National d’Histoire Naturelle (Paris), the Museo Nacional de Ciencias Naturales (Madrid), the Museo Argentino de Ciencias Naturales (Buenos Aires), the Museo de La Plata (Argentina), and the Naturhistorisches Museum Basel (Switzerland), for granting me access to the collections. We would also show our appreciation to the following organisations for partially funding this research: la Caixa; Deutscher Akademischer Austausch Dienst (DAAD); the University of Barcelona (UB); Agència de Gestió d’Ajuts Universitaris i de Recerca (AGAUR); Departament d’Innovació, Universitats i Empresa de la Generalitat de Catalunya; and the European Social Fund (ESF). Finally, this work was completed with the assistance of funds from research grants CGL2005-04402/BOS and CGL2008-00832/BOS from the former Ministerio de Educación y Ciencia (MEC) of Spain.

