## Supplementary Tables for "Scaling pattern of the carnivoran hind limb: Main deviations from a conservative pattern"

**Supplementary Material**

**Table S1. Branch length transformations used for phylogenetically independent contrasts.** Variable names are listed in Table 3. Abbreviations: exp., exponential transformation; Gra., transformation of Grafen; ln, natural logarithm transformation; Nee, transformation of Nee; $\rho_x$, Grafen's rho transform, where x indicates the value of rho; untr., untransformed branch lengths.

| | | | **phylogeny** | | | | | | | **locomotor type** | | | | | | |
|---|---|---|---|---|---|---|---|---|---|---|---|---|---|---|---|---|
| | **whole sample** | **fissipeds** | **Canidae** | **Mustelidae** | **Procyonidae** | **Ursidae** | **Felidae** | **Herpestidae** | **Viverridae** | **arboreal** | **semiarboreal** | **scansorial** | **terrestrial** | **semifossorial** | **semiaquatic** | **aquatic** |
| $L_f$ | $\rho_{0.5}$ | $\rho_{0.5}$ | ln | $\rho_{0.5}$ | untr. | $\rho_{0.1}$ | $\rho_{0.5}$ | $\rho_{0.5}$ | $\rho_{1.7}$ | exp. | $\rho_{0.8}$ | Nee | ln | $\rho_{0.3}$ | Nee | exp. |
| N | $\rho_{0.5}$ | $\rho_{0.5}$ | $\rho_{0.5}$ | $\rho_{0.5}$ | untr. | $\rho_{0.5}$ | $\rho_{0.5}$ | ln | Nee | $\rho_{0.1}$ | $\rho_{0.8}$ | Nee | ln | Nee | Nee | Nee |
| $d_{sf}$ | $\rho_{0.5}$ | $\rho_{0.5}$ | $\rho_{0.5}$ | $\rho_{0.5}$ | untr. | $\rho_{0.1}$ | $\rho_{0.5}$ | ln | $\rho_{1.7}$ | $\rho_{0.1}$ | $\rho_{0.8}$ | Nee | ln | untr. | Nee | Nee |
| $d_{tf}$ | $\rho_{0.5}$ | $\rho_{0.5}$ | ln | ln | Nee | $\rho_{0.5}$ | $\rho_{0.5}$ | ln | $\rho_{1.7}$ | $\rho_{0.1}$ | $\rho_{0.8}$ | Nee | ln | untr. | Nee | ln |
| $\varphi$ | exp. | exp. | $\rho_{0.5}$ | $\rho_{0.5}$ | untr. | $\rho_{0.1}$ | $\rho_{0.5}$ | untr. | exp. | $\rho_{0.1}$ | exp. | exp. | exp. | $\rho_{0.3}$ | Nee | $\rho_{0.8}$ |
| FR | $\rho_{0.5}$ | exp. | ln | exp. | ln | $\rho_{0.5}$ | $\rho_{0.5}$ | $\rho_{0.5}$ | Nee | exp. | Nee | Nee | $\rho_{0.5}$ | $\rho_{0.3}$ | Nee | $\rho_{0.8}$ |
| $L_t$ | Nee | $\rho_{0.5}$ | ln | $\rho_{0.5}$ | untr. | $\rho_{0.1}$ | $\rho_{0.5}$ | untr. | Nee | $\rho_{0.1}$ | $\rho_{0.8}$ | Nee | ln | $\rho_{0.3}$ | exp. | $\rho_{0.8}$ |
| $d_{st}$ | Nee | Nee | Nee | $\rho_{0.5}$ | untr. | $\rho_{0.5}$ | $\rho_{0.5}$ | ln | $\rho_{1.7}$ | $\rho_{0.1}$ | $\rho_{0.8}$ | Nee | ln | $\rho_{0.3}$ | Nee | Nee |
| $d_{tt}$ | Nee | Nee | $\rho_{0.5}$ | $\rho_{0.5}$ | untr. | $\rho_{0.5}$ | $\rho_{0.5}$ | untr. | $\rho_{1.7}$ | $\rho_{0.1}$ | ln | Nee | Nee | $\rho_{0.3}$ | Nee | ln |
| TR | $\rho_{0.5}$ | $\rho_{0.5}$ | Nee | $\rho_{0.5}$ | ln | $\rho_{0.5}$ | $\rho_{0.5}$ | $\rho_{0.5}$ | Nee | $\rho_{0.1}$ | ln | Nee | Nee | $\rho_{0.3}$ | Nee | Nee |
| $L_c$ | Nee | $\rho_{0.5}$ | Nee | ln | | | untr. | | $\rho_{1.7}$ | exp. | | Nee | Nee | | Nee | |
| r | $\rho_{0.5}$ | $\rho_{0.5}$ | Nee | ln | | | untr. | | exp. | exp. | | Nee | Nee | | Nee | |
| $d_{sc}$ | $\rho_{0.5}$ | $\rho_{0.5}$ | ln | ln | | | untr. | | $\rho_{1.7}$ | exp. | | Nee | ln | | Nee | |
| $d_{tc}$ | Nee | $\rho_{0.5}$ | $\rho_{0.5}$ | ln | | | untr. | | $\rho_{1.7}$ | exp. | | Nee | ln | | Nee | |

### Tables SR1 to SR13 – Results of traditional and PIC regressions

As indicated in Table 3, the following tables present the regression results for each variable. Both the results using traditional regression methods and phylogenetically independent contrasts (PIC) are shown for the whole sample and each of the subsamples (fissipeds, by Family, and by locomotor type). In each case, it is indicated (in the "sim." columns) whether the theoretical values proposed by the geometric similarity hypothesis (G), the elastic similarity hypothesis (E), or both (B), are included in the 95% confidence interval for the slope *b* (95% $CI_b$). Furthermore, when neither theoretical value is included in the 95% $CI_b$, it is indicated whether there is positive allometry (+; *b* is higher than both theoretical values), negative allometry (–; *b* is lower than both theoretical values), or both (nei.; *b* is higher than one theoretical value and lower than the other). Finally, the results of the comparison between the allometric coefficients obtained with each methodology are presented in the last column ($b_{trad} \neq b_{PIC}$): a cross (×) indicates no significant differences, while a tick (✓) denotes that the slopes are significantly different from each other ($p < 0.05$).

Variable names and abbreviations are given in Table 3, while the following abbreviations are common to all following tables: 95% $CI_a$, 95% confidence interval for the coefficient *a*; 95% $CI_b$, 95% confidence interval for the allometric coefficient *b*; n, sample size; n.s., unable to test differences due to non-significant regression; R, correlation coefficient; sim., similarity. Results in ***grey italics*** denote non-significant regressions.

| **SR1 – $L_f$** | traditional regression | | | | | | | PIC regression | | | | | |
|---|---|---|---|---|---|---|---|---|---|---|---|---|---|
| | **n** | ***a*** | **95% CI$_a$** | **$b_{trad}$** | **95% CI$_b$** | **R** | **sim.** | **n** | **$b_{PIC}$** | **95% CI$_b$** | **R** | **sim.** | **$b_{trad} \neq b_{PIC}$** |
| **whole sample** | 136 | 5.977 | 4.641 – 7.734 | 0.324 | 0.290 – 0.358 | 0.818 | G | 135 | 0.308 | 0.284 – 0.332 | 0.887 | G | × |
| **fissipeds** | 129 | 4.385 | 3.704 – 5.217 | 0.368 | 0.347 – 0.387 | 0.934 | + | 128 | 0.319 | 0.298 – 0.340 | 0.928 | G | ✓ |
| **Family** | | | | | | | | | | | | | |
| Canidae | 17 | 3.151 | 1.641 – 10.351 | 0.416 | 0.286 – 0.489 | 0.935 | G | 16 | 0.415 | 0.319 – 0.511 | 0.908 | G | × |
| Mustelidae | 32 | 6.192 | 4.118 – 9.310 | 0.296 | 0.244 – 0.348 | 0.882 | B | 31 | 0.297 | 0.248 – 0.346 | 0.894 | B | × |
| Procyonidae | 7 | 10.662 | 3.852 – 29.513 | 0.266 | 0.140 – 0.392 | 0.911 | B | 6 | 0.257 | 0.108 – 0.406 | 0.885 | B | × |
| Ursidae | 7 | 6.463 | 1.458 – 28.650 | 0.324 | 0.197 – 0.451 | 0.940 | B | 6 | 0.328 | 0.188 – 0.468 | 0.939 | B | × |
| Felidae | 26 | 9.953 | 8.163 – 12.134 | 0.295 | 0.274 – 0.316 | 0.986 | nei. | 25 | 0.307 | 0.279 – 0.335 | 0.977 | G | × |
| Herpestidae | 11 | 6.725 | 4.106 –11.016 | 0.310 | 0.241 – 0.378 | 0.956 | B | 10 | 0.304 | 0.233 – 0.375 | 0.953 | B | × |
| Eupleridae | 5 | 5.076 | 2.990 – 8.616 | 0.367 | 0.294 – 0.440 | 0.994 | G | | | | | | |
| Viverridae | 14 | 13.607 | 9.100 – 20.347 | 0.234 | 0.184 – 0.283 | 0.941 | E | 13 | 0.256 | 0.129 – 0.383 | 0.622 | B | × |
| **Locomotor type** | | | | | | | | | | | | | |
| arboreal | 7 | 8.163 | 2.193 – 30.900 | 0.301 | 0.138 – 0.464 | 0.883 | B | 6 | 0.335 | 0.326 – 0.344 | 1.000 | G | × |
| semiarboreal | 10 | 9.910 | 6.787 – 14.138 | 0.286 | 0.237 – 0.335 | 0.977 | B | 9 | 0.253 | 0.160 – 0.346 | 0.899 | B | × |
| scansorial | 45 | 7.244 | 6.084 – 8.894 | 0.322 | 0.303 – 0.341 | 0.982 | G | 44 | 0.322 | 0.299 – 0.345 | 0.972 | G | × |
| terrestrial | 48 | 3.469 | 2.694 – 4.897 | 0.395 | 0.353 – 0.425 | 0.964 | + | 47 | 0.340 | 0.299 – 0.381 | 0.915 | G | ✓ |
| semifossorial | 7 | 5.100 | 2.139 – 12.958 | 0.323 | 0.214 – 0.432 | 0.956 | B | 6 | 0.338 | 0.213 – 0.463 | 0.954 | B | × |
| semiaquatic | 11 | 9.210 | 3.391 – 25.068 | 0.243 | 0.126 – 0.360 | 0.769 | B | 10 | 0.253 | 0.149 – 0.357 | 0.844 | B | × |
| aquatic | ***8*** | ***13.106*** | ***1.968 – 87.560*** | ***0.168*** | ***0.010 – 0.327*** | ***0.330*** | ***E*** | 7 | 0.289 | 0.153 – 0.425 | 0.894 | B | n.s. |

| SR2 – N | traditional regression | | | | | | | PIC regression | | | | | |
|---|---|---|---|---|---|---|---|---|---|---|---|---|---|
| | n | *a* | 95% CI$_a$ | $b_{trad}$ | 95% CI$_b$ | R | sim. | n | $b_{PIC}$ | 95% CI$_b$ | R | sim. | $b_{trad} \neq b_{PIC}$ |
| **whole sample** | 136 | 0.699 | 0.571 – 0.860 | 0.335 | 0.309 – 0.362 | 0.945 | G | 135 | 0.344 | 0.322 – 0.366 | 0.929 | G | × |
| **fissipeds** | 129 | 0.541 | 0.483 – 0.610 | 0.369 | 0.355 – 0.383 | 0.977 | + | 128 | 0.353 | 0.332 – 0.374 | 0.943 | G | × |
| **Family** | | | | | | | | | | | | | |
| Canidae | 17 | 0.383 | 0.228 – 0.852 | 0.408 | 0.319 – 0.468 | 0.937 | G | 16 | 0.415 | 0.319 – 0.511 | 0.908 | G | × |
| Mustelidae | 32 | 0.604 | 0.460 – 0.791 | 0.347 | 0.313 – 0.381 | 0.964 | G | 31 | 0.344 | 0.296 – 0.392 | 0.928 | G | × |
| Procyonidae | 7 | 0.592 | 0.188 – 1.864 | 0.368 | 0.226 – 0.510 | 0.942 | B | 6 | 0.362 | 0.183 – 0.541 | 0.917 | B | × |
| Ursidae | 7 | 2.879 | 0.441 – 18.788 | 0.235 | 0.075 – 0.395 | 0.807 | B | ***6*** | ***0.236*** | ***0.065 – 0.407*** | ***0.811*** | ***B*** | n.s. |
| Felidae | 26 | 0.694 | 0.563 – 0.854 | 0.340 | 0.318 – 0.362 | 0.988 | G | 25 | 0.336 | 0.304 – 0.368 | 0.975 | G | × |
| Herpestidae | 11 | 0.770 | 0.480 – 1.233 | 0.328 | 0.262 – 0.394 | 0.964 | G | 10 | 0.318 | 0.245 – 0.391 | 0.955 | B | × |
| Eupleridae | 5 | 0.410 | 0.285 – 0.589 | 0.413 | 0.363 – 0.463 | 0.998 | + | | | | | | |
| Viverridae | 14 | 0.719 | 0.341 – 1.514 | 0.331 | 0.239 – 0.423 | 0.897 | B | 13 | 0.367 | 0.258 – 0.476 | 0.885 | G | × |
| **Locomotor type** | | | | | | | | | | | | | |
| arboreal | 7 | 0.390 | 0.205 – 0.744 | 0.412 | 0.332 – 0.492 | 0.986 | G | 6 | 0.414 | 0.325 – 0.503 | 0.985 | G | × |
| semiarboreal | 10 | 0.915 | 0.561 – 1.492 | 0.310 | 0.246 – 0.373 | 0.968 | B | 9 | 0.279 | 0.196 – 0.362 | 0.934 | B | × |
| scansorial | 45 | 0.533 | 0.443 – 0.641 | 0.373 | 0.353 – 0.393 | 0.984 | + | 44 | 0.359 | 0.329 – 0.389 | 0.961 | G | × |
| terrestrial | 48 | 0.509 | 0.425 – 0.638 | 0.375 | 0.350 – 0.397 | 0.974 | + | 47 | 0.350 | 0.310 – 0.390 | 0.923 | G | × |
| semifossorial | 7 | 0.639 | 0.282 – 1.450 | 0.352 | 0.249 – 0.455 | 0.967 | B | 6 | 0.372 | 0.214 – 0.530 | 0.940 | B | × |
| semiaquatic | 11 | 0.774 | 0.429 – 1.397 | 0.315 | 0.246 – 0.384 | 0.957 | B | 10 | 0.340 | 0.253 – 0.427 | 0.943 | G | × |
| aquatic | ***8*** | ***0.646*** | ***0.041 – 10.165*** | ***0.297*** | ***0.066 – 0.528*** | ***0.627*** | ***B*** | ***7*** | ***0.296*** | ***0.064 – 0.528*** | ***0.666*** | ***B*** | n.s. |

| **SR3 – $d_{sf}$** | **traditional regression** | | | | | | | **PIC regression** | | | | | |
|---|---|---|---|---|---|---|---|---|---|---|---|---|---|
| | **n** | ***a*** | **95% CI$_a$** | **$b_{trad}$** | **95% CI$_b$** | **R** | **sim.** | **n** | **$b_{PIC}$** | **95% CI$_b$** | **R** | **sim.** | **$b_{trad} \neq b_{PIC}$** |
| **whole sample** | 136 | 0.505 | 0.441 – 0.583 | 0.323 | 0.306 – 0.339 | 0.961 | G | 135 | 0.332 | 0.312 – 0.352 | 0.935 | G | × |
| **fissipeds** | 129 | 0.433 | 0.388 – 0.486 | 0.343 | 0.330 – 0.355 | 0.972 | G | 128 | 0.334 | 0.314 – 0.354 | 0.940 | G | × |
| **Family** | | | | | | | | | | | | | |
| Canidae | 17 | 0.228 | 0.133 – 0.593 | 0.412 | 0.305 – 0.472 | 0.951 | B | 16 | 0.401 | 0.324 – 0.478 | 0.938 | B | × |
| Mustelidae | 32 | 0.485 | 0.379 – 0.622 | 0.314 | 0.283 – 0.346 | 0.963 | G | 31 | 0.316 | 0.268 – 0.364 | 0.912 | G | × |
| Procyonidae | 7 | 0.601 | 0.295 – 1.223 | 0.315 | 0.227 – 0.403 | 0.970 | B | 6 | 0.310 | 0.195 – 0.425 | 0.954 | B | × |
| Ursidae | 7 | 0.524 | 0.090 – 3.062 | 0.324 | 0.174 – 0.474 | 0.915 | B | 6 | 0.325 | 0.160 – 0.490 | 0.912 | B | × |
| Felidae | 26 | 0.514 | 0.438 – 0.603 | 0.331 | 0.314 – 0.348 | 0.993 | G | 25 | 0.332 | 0.306 – 0.358 | 0.983 | G | × |
| Herpestidae | 11 | 0.576 | 0.364 – 0.912 | 0.314 | 0.249 – 0.378 | 0.963 | B | 10 | 0.311 | 0.250 – 0.372 | 0.967 | G | × |
| Eupleridae | 5 | 0.229 | 0.100 – 0.525 | 0.451 | 0.337 – 0.565 | 0.990 | E | | | | | | |
| Viverridae | 14 | 0.562 | 0.311 – 1.019 | 0.313 | 0.240 – 0.387 | 0.928 | B | 13 | 0.364 | 0.211 – 0.517 | 0.749 | B | × |
| **Locomotor type** | | | | | | | | | | | | | |
| arboreal | 7 | 0.470 | 0.190 – 1.163 | 0.342 | 0.230 – 0.455 | 0.958 | B | 6 | 0.350 | 0.229 – 0.471 | 0.960 | B | × |
| semiarboreal | 10 | 0.484 | 0.253 – 0.927 | 0.343 | 0.258 – 0.427 | 0.953 | B | 9 | 0.335 | 0.265 – 0.405 | 0.969 | B | × |
| scansorial | 45 | 0.484 | 0.424 – 0.552 | 0.335 | 0.321 – 0.349 | 0.990 | G | 44 | 0.338 | 0.314 – 0.362 | 0.973 | G | × |
| terrestrial | 48 | 0.409 | 0.346 – 0.511 | 0.346 | 0.321 – 0.365 | 0.969 | G | 47 | 0.330 | 0.286 – 0.374 | 0.896 | G | × |
| semifossorial | 7 | 0.653 | 0.328 – 1.301 | 0.282 | 0.195 – 0.368 | 0.963 | G | 6 | 0.306 | 0.166 – 0.446 | 0.930 | B | × |
| semiaquatic | 11 | 0.581 | 0.298 – 1.134 | 0.294 | 0.216 – 0.372 | 0.936 | G | 10 | 0.314 | 0.228 – 0.400 | 0.934 | B | × |
| aquatic | 8 | 0.133 | 0.004 – 4.009 | 0.410 | 0.125 – 0.695 | 0.718 | B | *7* | ***0.392*** | ***0.137 – 0.647*** | ***0.784*** | ***B*** | n.s. |

| **SR4 – $d_{tf}$** | **traditional regression** | | | | | | | **PIC regression** | | | | | |
|---|---|---|---|---|---|---|---|---|---|---|---|---|---|
| | **n** | ***a*** | **95% CI$_a$** | **$b_{trad}$** | **95% CI$_b$** | **R** | **sim.** | **n** | **$b_{PIC}$** | **95% CI$_b$** | **R** | **sim.** | **$b_{trad} \neq b_{PIC}$** |
| **whole sample** | 136 | 0.417 | 0.361 – 0.480 | 0.356 | 0.340 – 0.372 | 0.972 | nei. | 135 | 0.352 | 0.330 – 0.374 | 0.932 | G | × |
| **fissipeds** | 129 | 0.421 | 0.362 – 0.488 | 0.355 | 0.339 – 0.371 | 0.967 | nei. | 128 | 0.347 | 0.325 – 0.369 | 0.932 | G | × |
| **Family** | | | | | | | | | | | | | |
| Canidae | 17 | 0.247 | 0.126 – 0.724 | 0.405 | 0.283 – 0.479 | 0.952 | B | 16 | 0.402 | 0.322 – 0.482 | 0.934 | B | × |
| Mustelidae | 32 | 0.377 | 0.287 – 0.496 | 0.355 | 0.320 – 0.390 | 0.965 | B | 31 | 0.361 | 0.305 – 0.417 | 0.910 | G | × |
| Procyonidae | 7 | 0.831 | 0.399 – 1.729 | 0.285 | 0.194 – 0.376 | 0.961 | B | 6 | 0.279 | 0.191 – 0.367 | 0.967 | B | × |
| Ursidae | *7* | *1.691* | *0.193 – 14.835* | *0.240* | *0.055 – 0.425* | *0.743* | *B* | *6* | *0.241* | *0.043 – 0.439* | *0.748* | *B* | n.s. |
| Felidae | 26 | 0.494 | 0.375 – 0.652 | 0.342 | 0.312 – 0.371 | 0.979 | G | 25 | 0.343 | 0.293 – 0.393 | 0.939 | G | × |
| Herpestidae | 11 | 0.812 | 0.472 – 1.398 | 0.277 | 0.202 – 0.353 | 0.932 | G | 10 | 0.264 | 0.183 – 0.345 | 0.918 | G | × |
| Eupleridae | 5 | 0.605 | 0.449 – 0.815 | 0.330 | 0.288 – 0.371 | 0.998 | G | | | | | | |
| Viverridae | 14 | 0.733 | 0.409 – 1.314 | 0.292 | 0.220 – 0.364 | 0.919 | G | 13 | 0.301 | 0.208 – 0.394 | 0.874 | B | × |
| **Locomotor type** | | | | | | | | | | | | | |
| arboreal | 7 | 0.522 | 0.212 – 1.285 | 0.339 | 0.228 – 0.451 | 0.958 | B | 6 | 0.342 | 0.219 – 0.465 | 0.957 | B | × |
| semiarboreal | 10 | 0.603 | 0.336 – 1.085 | 0.320 | 0.244 – 0.397 | 0.956 | B | 9 | 0.305 | 0.232 – 0.378 | 0.958 | B | × |
| scansorial | 45 | 0.470 | 0.390 – 0.566 | 0.349 | 0.328 – 0.369 | 0.982 | G | 44 | 0.339 | 0.304 – 0.374 | 0.944 | G | × |
| terrestrial | 48 | 0.418 | 0.341 – 0.526 | 0.348 | 0.323 – 0.369 | 0.965 | G | 47 | 0.339 | 0.300 – 0.378 | 0.921 | B | × |
| semifossorial | 7 | 0.612 | 0.284 – 1.320 | 0.299 | 0.202 – 0.396 | 0.960 | B | 6 | 0.326 | 0.187 – 0.465 | 0.939 | B | × |
| semiaquatic | 11 | 0.486 | 0.224 – 1.057 | 0.331 | 0.240 – 0.422 | 0.931 | B | 10 | 0.367 | 0.259 – 0.475 | 0.923 | B | × |
| aquatic | 8 | 0.092 | 0.006 – 1.479 | 0.482 | 0.249 – 0.715 | 0.875 | B | 7 | 0.475 | 0.206 – 0.744 | 0.842 | B | × |

| SR5 – FR | traditional regression | | | | | | | PIC regression | | | | | |
|---|---|---|---|---|---|---|---|---|---|---|---|---|---|
| | **n** | ***a*** | **95% CI$_a$** | **$b_{trad}$** | **95% CI$_b$** | **R** | **sim.** | **n** | **$b_{PIC}$** | **95% CI$_b$** | **R** | **sim.** | **$b_{trad} \neq b_{PIC}$** |
| **whole sample** | 136 | 0.024 | 0.018 – 0.033 | 0.142 | 0.108 – 0.177 | 0.318 | E | 135 | 0.131 | 0.110 – 0.152 | 0.289 | E | × |
| **fissipeds** | *129* | *0.182* | *0.153 – 1.056* | *-0.097* | *-0.305 – -0.076* | *0.104* | *–* | *128* | *-0.135* | *-0.159 – -0.111* | *0.001* | *–* | n.s. |
| **Family** | | | | | | | | | | | | | |
| Canidae | *17* | *0.027* | *0.002 – 0.048* | *0.106* | *0.039 – 0.394* | *0.030* | *E* | *16* | *-0.129* | *-0.199 – -0.059* | *0.175* | *–* | n.s. |
| Mustelidae | *32* | *0.033* | *0.005 – 0.049* | *0.129* | *0.084 – 0.368* | *0.321* | *E* | *31* | *-0.151* | *-0.206 – -0.096* | *0.249* | *–* | n.s. |
| Procyonidae | *7* | *0.041* | *0.004 – 0.053* | *0.088* | *0.054 – 0.377* | *0.721* | *E* | 6 | 0.087 | 0.023 – 0.151 | 0.800 | E | n.s. |
| Ursidae | *7* | *0.175* | *0.080 – 2.071* | *-0.066* | *-0.275 – 0.006* | *0.149* | *B* | *6* | *-0.061* | *-0.140 – 0.006* | *0.237* | *B* | n.s. |
| Felidae | 26 | 0.040 | 0.033 – 0.050 | 0.064 | 0.040 – 0.081 | 0.587 | nei. | *25* | *0.076* | *0.046 – 0.106* | *0.346* | *nei.* | n.s. |
| Herpestidae | *11* | *0.032* | *0.003 – 0.079* | *0.140* | *0.010 – 0.483* | *0.045* | *E* | *10* | *0.139* | *0.032 – 0.246* | *0.086* | *E* | n.s. |
| Eupleridae | *5* | *0.040* | *0.007 – 0.130* | *0.101* | *-0.080 – 0.361* | *0.833* | *B* | | | | | | |
| Viverridae | 14 | 0.037 | 0.031 – 0.046 | 0.094 | 0.066 – 0.117 | 0.767 | nei. | 13 | 0.110 | 0.060 – 0.160 | 0.701 | E | × |
| **Locomotor type** | | | | | | | | | | | | | |
| arboreal | *7* | *0.033* | *0.001 – 0.096* | *0.112* | *-0.029 – 0.541* | *0.571* | *B* | 6 | 0.089 | 0.085 – 0.093 | 0.999 | E | n.s. |
| semiarboreal | *10* | *0.029* | *0.005 – 0.061* | *0.124* | *0.021 – 0.346* | *0.385* | *E* | *9* | *0.142* | *0.038 – 0.246* | *0.490* | *E* | n.s. |
| scansorial | *45* | *0.044* | *0.038 – 0.052* | *0.058* | *0.041 – 0.073* | *0.274* | *nei.* | *44* | *0.078* | *0.055 – 0.101* | *0.203* | *nei.* | n.s. |
| terrestrial | 48 | 0.165 | 0.130 – 0.197 | -0.090 | -0.112 – -0.060 | 0.508 | – | *47* | *-0.103* | *-0.133 – -0.073* | *0.192* | *–* | n.s. |
| semifossorial | *7* | *0.147* | *0.116 – 0.298* | *-0.059* | *-0.146 – -0.025* | *0.641* | *–* | *6* | *-0.066* | *-0.124 – -0.008* | *0.702* | *–* | n.s. |
| semiaquatic | 11 | 0.036 | 0.025 – 0.078 | 0.119 | 0.031 – 0.157 | 0.750 | E | 10 | 0.130 | 0.051 – 0.209 | 0.616 | E | × |
| aquatic | *8* | *0.003* | *$1.95 \cdot 10^{-4}$ – 0.401* | *0.334* | *-0.044 – 0.582* | *0.718* | *B* | *7* | *0.306* | *0.070 – 0.542* | *0.676* | *E* | n.s. |

| **SR6 – $L_t$** | **traditional regression** | | | | | | | **PIC regression** | | | | | |
|---|---|---|---|---|---|---|---|---|---|---|---|---|---|
| | **n** | ***a*** | **95% CI$_a$** | **$b_{trad}$** | **95% CI$_b$** | **R** | **sim.** | **n** | **$b_{PIC}$** | **95% CI$_b$** | **R** | **sim.** | **$b_{trad} \neq b_{PIC}$** |
| **whole sample** | 136 | 7.202 | 5.931 – 8.798 | 0.308 | 0.284 – 0.330 | 0.915 | nei. | 135 | 0.306 | 0.283 – 0.329 | 0.897 | nei. | × |
| **fissipeds** | 129 | 6.122 | 4.999 – 7.628 | 0.329 | 0.303 – 0.353 | 0.911 | G | 128 | 0.297 | 0.274 – 0.320 | 0.901 | nei. | ✓ |
| **Family** | | | | | | | | | | | | | |
| Canidae | 17 | 3.834 | 1.530 – 17.140 | 0.400 | 0.239 – 0.503 | 0.873 | B | 16 | 0.436 | 0.307 – 0.565 | 0.847 | G | × |
| Mustelidae | 32 | 6.849 | 4.608 – 10.179 | 0.290 | 0.240 – 0.341 | 0.885 | B | 31 | 0.282 | 0.233 – 0.331 | 0.885 | E | × |
| Procyonidae | 7 | 12.561 | 4.110 – 38.393 | 0.246 | 0.108 – 0.385 | 0.872 | B | 6 | 0.224 | 0.104 – 0.344 | 0.902 | B | × |
| Ursidae | 7 | 5.304 | 1.425 – 19.739 | 0.317 | 0.205 – 0.429 | 0.952 | B | 6 | 0.329 | 0.204 – 0.454 | 0.952 | B | × |
| Felidae | 26 | 14.249 | 10.755 – 18.878 | 0.255 | 0.225 – 0.285 | 0.961 | E | 25 | 0.280 | 0.241 – 0.319 | 0.945 | E | × |
| Herpestidae | 11 | 6.902 | 3.537 – 13.470 | 0.312 | 0.219 – 0.406 | 0.918 | B | 10 | 0.298 | 0.209 – 0.387 | 0.922 | B | × |
| Eupleridae | 5 | 8.448 | 2.701 – 26.422 | 0.307 | 0.150 – 0.464 | 0.960 | B | | | | | | |
| Viverridae | 14 | 18.548 | 13.015 – 26.433 | 0.196 | 0.153 – 0.240 | 0.935 | – | 13 | 0.240 | 0.181 – 0.299 | 0.922 | E | × |
| **Locomotor type** | | | | | | | | | | | | | |
| arboreal | 7 | 8.552 | 1.515 – 48.267 | 0.293 | 0.079 – 0.508 | 0.772 | B | *6* | *0.297* | *0.068 – 0.526* | *0.783* | *B* | n.s. |
| semiarboreal | 10 | 15.163 | 9.625 – 23.888 | 0.232 | 0.173 – 0.291 | 0.950 | E | 9 | 0.201 | 0.108 – 0.294 | 0.832 | E | × |
| scansorial | 45 | 11.576 | 9.164 – 14.622 | 0.267 | 0.242 – 0.292 | 0.951 | E | 44 | 0.287 | 0.259 – 0.315 | 0.949 | nei. | × |
| terrestrial | 48 | 4.287 | 3.100 – 6.433 | 0.372 | 0.321 – 0.412 | 0.934 | G | 47 | 0.318 | 0.272 – 0.364 | 0.871 | G | ✓ |
| semifossorial | 7 | 9.724 | 3.727 – 25.372 | 0.240 | 0.120 – 0.361 | 0.899 | B | 6 | 0.258 | 0.115 – 0.401 | 0.896 | B | × |
| semiaquatic | 11 | 8.964 | 3.880 – 20.710 | 0.258 | 0.160 – 0.356 | 0.864 | B | 10 | 0.282 | 0.203 – 0.361 | 0.931 | B | × |
| aquatic | 8 | 6.817 | 3.210 – 14.474 | 0.287 | 0.224 – 0.350 | 0.976 | B | 7 | 0.258 | 0.185 – 0.331 | 0.963 | E | × |

| **SR7 – $d_{st}$** | traditional regression | | | | | | | PIC regression | | | | | |
|---|---|---|---|---|---|---|---|---|---|---|---|---|---|
| | **n** | ***a*** | **95% $CI_a$** | **$b_{trad}$** | **95% $CI_b$** | **R** | **sim.** | **n** | **$b_{PIC}$** | **95% $CI_b$** | **R** | **sim.** | **$b_{trad} \neq b_{PIC}$** |
| **whole sample** | 136 | 0.491 | 0.409 – 0.593 | 0.332 | 0.309 – 0.355 | 0.952 | G | 135 | 0.342 | 0.319 – 0.365 | 0.921 | G | × |
| **fissipeds** | 129 | 0.400 | 0.353 – 0.460 | 0.359 | 0.344 – 0.373 | 0.973 | nei. | 128 | 0.344 | 0.324 – 0.364 | 0.944 | G | × |
| **Family** | | | | | | | | | | | | | |
| Canidae | 17 | 0.246 | 0.159 – 0.511 | 0.408 | 0.327 – 0.454 | 0.968 | B | 16 | 0.385 | 0.321 – 0.449 | 0.954 | B | × |
| Mustelidae | 32 | 0.403 | 0.302 – 0.537 | 0.346 | 0.310 – 0.383 | 0.959 | B | 31 | 0.346 | 0.299 – 0.393 | 0.931 | B | × |
| Procyonidae | 7 | 0.888 | 0.256 – 3.077 | 0.267 | 0.113 – 0.420 | 0.865 | B | *6* | *0.255* | *0.082 – 0.428* | *0.839* | *B* | n.s. |
| Ursidae | 7 | 0.239 | 0.015 – 3.912 | 0.400 | 0.162 – 0.638 | 0.856 | B | *6* | *0.409* | *0.145 – 0.673* | *0.854* | *B* | n.s. |
| Felidae | 26 | 0.417 | 0.320 – 0.543 | 0.362 | 0.334 – 0.390 | 0.983 | E | 25 | 0.349 | 0.311 – 0.387 | 0.966 | B | × |
| Herpestidae | 11 | 0.885 | 0.572 – 1.370 | 0.263 | 0.202 – 0.324 | 0.951 | – | 10 | 0.262 | 0.198 – 0.326 | 0.948 | – | × |
| Eupleridae | 5 | 0.309 | 0.159 – 0.601 | 0.412 | 0.321 – 0.504 | 0.993 | B | | | | | | |
| Viverridae | 14 | 0.731 | 0.510 – 1.048 | 0.284 | 0.240 – 0.329 | 0.968 | – | 13 | 0.311 | 0.253 – 0.369 | 0.956 | G | × |
| **Locomotor type** | | | | | | | | | | | | | |
| arboreal | 7 | 0.475 | 0.222 – 1.019 | 0.342 | 0.248 – 0.437 | 0.971 | B | 6 | 0.343 | 0.243 – 0.443 | 0.972 | B | × |
| semiarboreal | 10 | 0.329 | 0.201 – 0.538 | 0.396 | 0.332 – 0.460 | 0.980 | B | 9 | 0.370 | 0.311 – 0.429 | 0.982 | E | × |
| scansorial | 45 | 0.444 | 0.370 – 0.532 | 0.353 | 0.333 – 0.372 | 0.983 | G | 44 | 0.352 | 0.316 – 0.388 | 0.942 | B | × |
| terrestrial | 48 | 0.407 | 0.339 – 0.514 | 0.353 | 0.329 – 0.373 | 0.970 | G | 47 | 0.328 | 0.294 – 0.362 | 0.938 | G | × |
| semifossorial | 7 | 0.471 | 0.222 – 1.001 | 0.333 | 0.238 – 0.428 | 0.969 | B | 6 | 0.342 | 0.233 – 0.451 | 0.967 | B | × |
| semiaquatic | 11 | 0.410 | 0.176 – 0.954 | 0.344 | 0.245 – 0.442 | 0.924 | B | 10 | 0.383 | 0.242 – 0.524 | 0.877 | B | × |
| aquatic | *8* | *0.261* | *0.007 – 10.243* | *0.351* | *0.044 – 0.659* | *0.483* | *B* | *7* | *0.377* | *0.027 – 0.727* | *0.466* | *B* | n.s. |

| **SR8 – $d_{tt}$** | **traditional regression** | | | | | | | **PIC regression** | | | | | |
|---|---|---|---|---|---|---|---|---|---|---|---|---|---|
| | **n** | ***a*** | **95% $CI_a$** | **$b_{trad}$** | **95% $CI_b$** | **R** | **sim.** | **n** | **$b_{PIC}$** | **95% $CI_b$** | **R** | **sim.** | **$b_{trad} \neq b_{PIC}$** |
| **whole sample** | 136 | 0.334 | 0.286 – 0.392 | 0.357 | 0.339 – 0.374 | 0.964 | nei. | 135 | 0.351 | 0.329 – 0.373 | 0.934 | G | × |
| **fissipeds** | 129 | 0.309 | 0.262 – 0.366 | 0.367 | 0.348 – 0.385 | 0.961 | E | 128 | 0.350 | 0.328 – 0.372 | 0.934 | G | × |
| **Family** | | | | | | | | | | | | | |
| Canidae | 17 | 0.212 | 0.119 – 0.591 | 0.418 | 0.306 – 0.481 | 0.947 | B | 16 | 0.421 | 0.338 – 0.504 | 0.934 | E | × |
| Mustelidae | 32 | 0.357 | 0.270 – 0.472 | 0.330 | 0.294 – 0.365 | 0.957 | G | 31 | 0.343 | 0.297 – 0.389 | 0.933 | B | × |
| Procyonidae | 7 | 0.361 | 0.139 – 0.942 | 0.347 | 0.228 – 0.465 | 0.955 | B | 6 | 0.340 | 0.190 – 0.490 | 0.935 | B | × |
| Ursidae | 7 | 0.552 | 0.049 – 6.234 | 0.308 | 0.101 – 0.514 | 0.812 | B | *6* | *0.309* | *0.093 – 0.525* | *0.826* | *B* | n.s. |
| Felidae | 26 | 0.408 | 0.316 – 0.526 | 0.348 | 0.321 – 0.375 | 0.983 | B | 25 | 0.362 | 0.319 – 0.405 | 0.960 | B | × |
| Herpestidae | 11 | 0.690 | 0.353 – 1.350 | 0.272 | 0.179 – 0.366 | 0.890 | G | 10 | 0.237 | 0.148 – 0.326 | 0.873 | – | × |
| Eupleridae | 5 | 0.357 | 0.080 – 1.597 | 0.364 | 0.158 – 0.571 | 0.951 | B | | | | | | |
| Viverridae | 14 | 0.493 | 0.367 – 0.661 | 0.308 | 0.271 – 0.344 | 0.982 | G | 13 | 0.324 | 0.269 – 0.379 | 0.964 | B | × |
| **Locomotor type** | | | | | | | | | | | | | |
| arboreal | 7 | 0.318 | 0.105 – 0.961 | 0.362 | 0.225 – 0.499 | 0.944 | B | 6 | 0.361 | 0.217 – 0.505 | 0.947 | B | × |
| semiarboreal | 10 | 0.506 | 0.307 – 0.834 | 0.311 | 0.246 – 0.376 | 0.967 | B | 9 | 0.313 | 0.238 – 0.388 | 0.958 | B | × |
| scansorial | 45 | 0.374 | 0.310 – 0.453 | 0.352 | 0.331 – 0.372 | 0.982 | G | 44 | 0.357 | 0.324 – 0.390 | 0.954 | B | × |
| terrestrial | 48 | 0.291 | 0.218 – 0.431 | 0.375 | 0.330 – 0.409 | 0.954 | B | 47 | 0.351 | 0.310 – 0.392 | 0.921 | B | × |
| semifossorial | 7 | 0.380 | 0.148 – 0.981 | 0.314 | 0.195 – 0.434 | 0.944 | B | 6 | 0.318 | 0.190 – 0.446 | 0.947 | B | × |
| semiaquatic | 11 | 0.433 | 0.226 – 0.831 | 0.311 | 0.234 – 0.387 | 0.945 | B | 10 | 0.354 | 0.255 – 0.453 | 0.931 | B | × |
| aquatic | 8 | 0.063 | 0.005 – 0.765 | 0.483 | 0.273 – 0.692 | 0.900 | B | 7 | 0.446 | 0.248 – 0.644 | 0.905 | B | × |

| SR9 – TR | traditional regression | | | | | | | PIC regression | | | | | |
|---|---|---|---|---|---|---|---|---|---|---|---|---|---|
| | n | *a* | 95% CI$_a$ | $b_{trad}$ | 95% CI$_b$ | R | sim. | n | $b_{PIC}$ | 95% CI$_b$ | R | sim. | $b_{trad} \neq b_{PIC}$ |
| **whole sample** | 136 | 0.031 | 0.027 – 0.036 | 0.114 | 0.097 – 0.128 | 0.297 | E | 135 | 0.141 | 0.119 – 0.163 | 0.393 | E | ✓ |
| **fissipeds** | 129 | 0.029 | 0.025 – 0.035 | 0.124 | 0.105 – 0.140 | 0.398 | E | 128 | 0.134 | 0.113 – 0.155 | 0.466 | E | × |
| **Family** | | | | | | | | | | | | | |
| Canidae | *17* | *0.014* | *$2.36 \cdot 10^{-4}$ – 0.028* | *0.176* | *0.093 – 0.629* | *0.259* | *E* | *16* | *-0.242* | *-0.372 – -0.112* | *0.251* | *–* | n.s. |
| Mustelidae | 32 | 0.035 | 0.025 – 0.053 | 0.124 | 0.075 – 0.161 | 0.603 | E | 31 | 0.117 | 0.083 – 0.151 | 0.621 | E | × |
| Procyonidae | *7* | *0.033* | *0.002 – 0.052* | *0.116* | *0.059 – 0.452* | *0.138* | *E* | *6* | *0.124* | *-0.028 – 0.276* | *0.121* | *B* | n.s. |
| Ursidae | *7* | *0.023* | *$2.14 \cdot 10^{-4}$ – 0.165* | *0.139* | *-0.031 – 0.538* | *0.272* | *B* | *6* | *0.156* | *-0.024 – 0.336* | *0.359* | *B* | n.s. |
| Felidae | 26 | 0.021 | 0.0.16 – 0.028 | 0.145 | 0.110 – 0.170 | 0.767 | E | 25 | 0.142 | 0.090 – 0.194 | 0.510 | E | × |
| Herpestidae | *11* | *0.257* | *0.140 – 3.761* | *-0.147* | *-0.526 – -0.067* | *0.254* | *–* | *10* | *-0.143* | *-0.250 – -0.036* | *0.229* | *–* | n.s. |
| Eupleridae | *5* | *0.022* | *$5.02 \cdot 10^{-4}$ – 0.046* | *0.173* | *0.091 – 0.718* | *0.664* | *E* | | | | | | |
| Viverridae | 14 | 0.031 | 0.026 – 0.046 | 0.116 | 0.068 – 0.142 | 0.787 | E | 13 | 0.115 | 0.065 – 0.165 | 0.735 | E | × |
| **Locomotor type** | | | | | | | | | | | | | |
| arboreal | *7* | *0.021* | *$1.05 \cdot 10^{-4}$ – 0.072* | *0.170* | *0.014 – 0.838* | *0.628* | *E* | 6 | 0.192 | 0.052 – 0.332 | 0.809 | E | n.s. |
| semiarboreal | 10 | 0.018 | 0.011 – 0.033 | 0.186 | 0.108 – 0.248 | 0.900 | E | 9 | 0.201 | 0.108 – 0.294 | 0.833 | E | × |
| scansorial | 45 | 0.027 | 0.022 – 0.034 | 0.125 | 0.101 – 0.147 | 0.744 | E | 44 | 0.134 | 0.097 – 0.171 | 0.442 | E | × |
| terrestrial | *48* | *0.206* | *0.165 – 0.646* | *-0.114* | *-0.357 – -0.086* | *0.046* | *–* | *47* | *0.132* | *0.093 – 0.171* | *0.126* | *E* | n.s. |
| semifossorial | 7 | 0.041 | 0.030 – 0.059 | 0.115 | 0.072 – 0.153 | 0.927 | E | 6 | 0.109 | 0.055 – 0.163 | 0.917 | E | × |
| semiaquatic | 11 | 0.027 | 0.015 – 0.062 | 0.146 | 0.051 – 0.212 | 0.654 | E | *10* | *0.190* | *0.065 – 0.315* | *0.508* | *E* | n.s. |
| aquatic | *8* | *3.323* | *0.080 – $4.71 \cdot 10^{5}$* | *-0.311* | *-1.298 – -0.018* | *0.357* | *–* | *7* | *-0.312* | *-0.630 – 0.006* | *0.240* | *G* | n.s. |

| SR10 – $L_c$ | traditional regression | | | | | | | PIC regression | | | | | |
|---|---|---|---|---|---|---|---|---|---|---|---|---|---|
| | n | *a* | 95% $CI_a$ | $b_{trad}$ | 95% $CI_b$ | R | sim. | n | $b_{PIC}$ | 95% $CI_b$ | R | sim. | $b_{trad} \neq b_{PIC}$ |
| **whole sample** | 85 | 1.409 | 1.098 – 1.828 | 0.342 | 0.311 – 0.370 | 0.954 | G | 84 | 0.338 | 0.313 – 0.363 | 0.939 | G | × |
| **fissipeds** | 84 | 1.280 | 1.049 – 1.638 | 0.354 | 0.325 – 0.377 | 0.961 | G | 83 | 0.339 | 0.314 – 0.364 | 0.943 | G | × |
| **Family** | | | | | | | | | | | | | |
| Canidae | 19 | 0.643 | 0.407 – 1.531 | 0.435 | 0.339 – 0.486 | 0.943 | + | 18 | 0.455 | 0.368 – 0.542 | 0.929 | + | × |
| Mustelidae | 15 | 1.256 | 0.794 – 1.987 | 0.340 | 0.281 – 0.400 | 0.957 | G | 14 | 0.296 | 0.235 – 0.357 | 0.941 | B | × |
| Procyonidae | | | | | | | | | | | | | |
| Ursidae | 5 | 0.412 | 0.005 – 33.229 | 0.425 | 0.059 – 0.792 | 0.883 | B | | | | | | |
| Felidae | 21 | 1.876 | 1.427 – 2.467 | 0.325 | 0.297 – 0.353 | 0.984 | G | 20 | 0.324 | 0.267 – 0.381 | 0.932 | G | × |
| Herpestidae | | | | | | | | | | | | | |
| Eupleridae | | | | | | | | | | | | | |
| Viverridae | 9 | 2.375 | 1.239 – 4.552 | 0.278 | 0.198 – 0.358 | 0.947 | B | 8 | 0.338 | 0.212 – 0.464 | 0.905 | B | × |
| **Locomotor type** | | | | | | | | | | | | | |
| arboreal | 6 | 1.171 | 0.399 – 3.437 | 0.363 | 0.232 – 0.494 | 0.966 | B | 5 | 0.417 | 0.407 – 0.427 | 1.000 | + | × |
| semiarboreal | 5 | 2.067 | 0.774 – 5.521 | 0.306 | 0.179 – 0.433 | 0.974 | B | | | | | | |
| scansorial | 35 | 1.697 | 1.272 – 2.238 | 0.326 | 0.294 – 0.358 | 0.963 | G | 33 | 0.318 | 0.286 – 0.350 | 0.960 | G | × |
| terrestrial | 30 | 1.023 | 0.715 – 1.912 | 0.378 | 0.304 – 0.418 | 0.961 | G | 30 | 0.357 | 0.311 – 0.403 | 0.941 | G | × |
| semifossorial | | | | | | | | | | | | | |
| semiaquatic | 6 | 1.103 | 0.175 – 6.945 | 0.359 | 0.133 – 0.585 | 0.891 | B | ***5*** | ***0.362*** | ***0.110 – 0.614*** | ***0.899*** | ***B*** | n.s. |
| aquatic | | | | | | | | | | | | | |

| **SR11 – r** | **traditional regression** | | | | | | | **PIC regression** | | | | | |
|---|---|---|---|---|---|---|---|---|---|---|---|---|---|
| | **n** | ***a*** | **95% CI$_a$** | **$b_{trad}$** | **95% CI$_b$** | **R** | **sim.** | **n** | **$b_{PIC}$** | **95% CI$_b$** | **R** | **sim.** | **$b_{trad} \neq b_{PIC}$** |
| **whole sample** | 85 | 0.997 | 0.806 – 1.264 | 0.350 | 0.322 – 0.376 | 0.964 | G | 84 | 0.349 | 0.322 – 0.376 | 0.938 | G | × |
| **fissipeds** | 84 | 0.914 | 0.770 – 1.135 | 0.361 | 0.336 – 0.381 | 0.969 | + | 83 | 0.359 | 0.332 – 0.386 | 0.939 | G | × |
| **Family** | | | | | | | | | | | | | |
| Canidae | 19 | 0.384 | 0.227 – 1.015 | 0.462 | 0.355 – 0.519 | 0.945 | + | 18 | 0.478 | 0.386 – 0.570 | 0.927 | + | × |
| Mustelidae | 15 | 0.904 | 0.579 – 1.409 | 0.350 | 0.293 – 0.408 | 0.962 | G | 14 | 0.305 | 0.244 – 0.366 | 0.946 | B | × |
| Procyonidae | | | | | | | | | | | | | |
| Ursidae | *5* | *0.253* | *0.001 – 45.479* | *0.449* | *0.016 – 0.882* | *0.851* | *B* | | | | | | |
| Felidae | 21 | 0.981 | 0.668 – 1.440 | 0.360 | 0.321 – 0.399 | 0.974 | G | 20 | 0.377 | 0.298 – 0.456 | 0.899 | G | × |
| Herpestidae | | | | | | | | | | | | | |
| Eupleridae | | | | | | | | | | | | | |
| Viverridae | 9 | 1.497 | 0.805 – 2.786 | 0.299 | 0.223 – 0.376 | 0.959 | B | 8 | 0.358 | 0.228 – 0.488 | 0.919 | B | × |
| **Locomotor type** | | | | | | | | | | | | | |
| arboreal | 6 | 0.751 | 0.172 – 3.283 | 0.382 | 0.203 – 0.562 | 0.941 | B | 5 | 0.417 | 0.407 – 0.427 | 1.000 | + | × |
| semiarboreal | 5 | 1.508 | 0.563 – 4.038 | 0.309 | 0.181 – 0.436 | 0.974 | B | | | | | | |
| scansorial | 35 | 1.102 | 0.856 – 1.425 | 0.342 | 0.312 – 0.370 | 0.972 | G | 33 | 0.339 | 0.303 – 0.375 | 0.955 | G | × |
| terrestrial | 30 | 0.741 | 0.544 – 1.316 | 0.385 | 0.318 – 0.420 | 0.965 | G | 30 | 0.369 | 0.322 – 0.416 | 0.942 | G | × |
| semifossorial | | | | | | | | | | | | | |
| semiaquatic | 6 | 0.873 | 0.169 – 4.522 | 0.358 | 0.156 – 0.560 | 0.913 | B | *5* | *0.355* | *0.127 – 0.583* | *0.914* | *B* | n.s. |
| aquatic | | | | | | | | | | | | | |

| SR12 – $d_{sc}$ | traditional regression | | | | | | | PIC regression | | | | | |
|---|---|---|---|---|---|---|---|---|---|---|---|---|---|
| | **n** | ***a*** | **95% CI$_a$** | **$b_{trad}$** | **95% CI$_b$** | **R** | **sim.** | **n** | **$b_{PIC}$** | **95% CI$_b$** | **R** | **sim.** | **$b_{trad} \neq b_{PIC}$** |
| **whole sample** | 85 | 0.495 | 0.391 – 0.636 | 0.346 | 0.317 – 0.373 | 0.961 | G | 84 | 0.360 | 0.329 – 0.391 | 0.919 | B | × |
| **fissipeds** | 84 | 0.452 | 0.377 – 0.556 | 0.357 | 0.334 – 0.377 | 0.967 | E | 83 | 0.372 | 0.340 – 0.404 | 0.922 | E | × |
| **Family** | | | | | | | | | | | | | |
| Canidae | 19 | 0.137 | 0.055 – 0.511 | 0.486 | 0.342 – 0.589 | 0.905 | E | 18 | 0.494 | 0.370 – 0.618 | 0.874 | E | × |
| Mustelidae | 15 | 0.452 | 0.278 – 0.734 | 0.353 | 0.290 – 0.416 | 0.955 | B | 14 | 0.325 | 0.255 – 0.395 | 0.934 | B | × |
| Procyonidae | | | | | | | | | | | | | |
| Ursidae | 5 | 0.070 | 0.001 – 9.631 | 0.505 | 0.094 – 0.915 | 0.896 | B | | | | | | |
| Felidae | 21 | 0.431 | 0.292 – 0.636 | 0.366 | 0.326 – 0.406 | 0.974 | B | 20 | 0.397 | 0.310 – 0.484 | 0.890 | B | × |
| Herpestidae | | | | | | | | | | | | | |
| Eupleridae | | | | | | | | | | | | | |
| Viverridae | 9 | 0.878 | 0.523 – 1.474 | 0.278 | 0.214 – 0.341 | 0.967 | G | 8 | 0.320 | 0.213 – 0.427 | 0.933 | B | × |
| **Locomotor type** | | | | | | | | | | | | | |
| arboreal | 6 | 0.544 | 0.119 – 2.494 | 0.336 | 0.141 – 0.511 | 0.912 | B | 5 | 0.286 | 0.278 – 0.294 | 1.000 | – | × |
| semiarboreal | 5 | 0.700 | 0.285 – 1.718 | 0.318 | 0.201 – 0.434 | 0.980 | B | | | | | | |
| scansorial | 35 | 0.557 | 0.444 – 0.715 | 0.337 | 0.311 – 0.362 | 0.981 | G | 33 | 0.338 | 0.303 – 0.373 | 0.956 | G | × |
| terrestrial | 30 | 0.364 | 0.256 – 0.640 | 0.379 | 0.316 – 0.418 | 0.954 | B | 30 | 0.378 | 0.318 – 0.438 | 0.909 | B | × |
| semifossorial | | | | | | | | | | | | | |
| semiaquatic | 6 | 0.402 | 0.080 – 2.005 | 0.372 | 0.174 – 0.569 | 0.924 | B | ***5*** | ***0.348*** | ***0.073 – 0.623*** | ***0.868*** | ***B*** | n.s. |
| aquatic | | | | | | | | | | | | | |

| SR13 – $d_{tc}$ | traditional regression | | | | | | | PIC regression | | | | | |
|---|---|---|---|---|---|---|---|---|---|---|---|---|---|
| | **n** | ***a*** | **95% CI$_a$** | **$b_{trad}$** | **95% CI$_b$** | **R** | **sim.** | **n** | **$b_{PIC}$** | **95% CI$_b$** | **R** | **sim.** | **$b_{trad} \neq b_{PIC}$** |
| **whole sample** | 85 | 0.241 | 0.194 – 0.306 | 0.346 | 0.319 – 0.370 | 0.952 | G | 84 | 0.358 | 0.322 – 0.394 | 0.890 | B | × |
| **fissipeds** | 84 | 0.236 | 0.185 – 0.309 | 0.348 | 0.318 – 0.377 | 0.948 | B | 83 | 0.351 | 0.314 – 0.388 | 0.876 | B | × |
| **Family** | | | | | | | | | | | | | |
| Canidae | 19 | 0.122 | 0.063 – 0.401 | 0.424 | 0.288 – 0.497 | 0.908 | B | 18 | 0.436 | 0.329 – 0.543 | 0.879 | B | × |
| Mustelidae | 15 | 0.239 | 0.136 – 0.419 | 0.339 | 0.266 – 0.411 | 0.934 | B | 14 | 0.324 | 0.227 – 0.421 | 0.870 | B | × |
| Procyonidae | | | | | | | | | | | | | |
| Ursidae | *5* | *0.001* | *0.000 – 476.235* | *0.763* | *-0.305 – 1.831* | *0.648* | *B* | | | | | | |
| Felidae | 21 | 0.243 | 0.154 – 0.385 | 0.353 | 0.306 – 0.400 | 0.961 | B | 20 | 0.332 | 0.256 – 0.408 | 0.879 | B | × |
| Herpestidae | | | | | | | | | | | | | |
| Eupleridae | | | | | | | | | | | | | |
| Viverridae | 9 | 0.393 | 0.212 – 0.728 | 0.289 | 0.213 – 0.364 | 0.956 | G | 8 | 0.335 | 0.225 – 0.445 | 0.935 | B | × |
| **Locomotor type** | | | | | | | | | | | | | |
| arboreal | 6 | 0.192 | 0.056 – 0.656 | 0.368 | 0.218 – 0.518 | 0.956 | B | 5 | 0.399 | 0.384 – 0.414 | 1.000 | + | × |
| semiarboreal | *5* | *0.319* | *0.036 – 2.806* | *0.314* | *0.033 – 0.596* | *0.873* | *B* | | | | | | |
| scansorial | 35 | 0.286 | 0.198 – 0.404 | 0.329 | 0.292 – 0.369 | 0.933 | G | 33 | 0.316 | 0.262 – 0.370 | 0.879 | G | × |
| terrestrial | 30 | 0.194 | 0.128 – 0.349 | 0.369 | 0.302 – 0.415 | 0.953 | B | 30 | 0.358 | 0.294 – 0.422 | 0.883 | B | × |
| semifossorial | | | | | | | | | | | | | |
| semiaquatic | 6 | 0.287 | 0.072 – 1.148 | 0.320 | 0.149 – 0.490 | 0.923 | B | *5* | *0.316* | *0.118 – 0.514* | *0.919* | *B* | n.s. |
| aquatic | | | | | | | | | | | | | |

**Tables SR14 to SR26. Results of the complex allometry test.**

In each case, it is indicated (in the "$D \neq 1$" column) whether the exponent of complex allometry ($D$) is significantly different from 1. Results in *grey italics* denote non-significant regressions. Variable names are listed in Table 3. Abbreviations: 95% $CI_C$, 95% confidence interval for the coefficient ($C$); 95% $CI_D$, 95% confidence interval for the exponent of complex allometry ($D$); 95% $CI_{ln\ A}$, 95% confidence interval for $ln\ A$; n, sample size; n.c., the model did not converge in a realistic solution; n.s., although the model did converge in a realistic solution, it was not significant according to the associated correlation coefficient (R).

| **SR14 – $L_f$** | **n** | ***ln A*** | **95% $CI_{ln\ A}$** | ***C*** | **95% $CI_C$** | ***D*** | **95% $CI_D$** | **R** | **$D \neq 1$** |
|---|---|---|---|---|---|---|---|---|---|
| **whole sample** | 136 | 5.365 | 5.198 – 5.532 | 0.032 | $-1.72 \cdot 10^{-4}$ – 0.065 | 1.999 | 1.514 – 2.484 | 0.853 | ✓ ($D$>1) |
| **fissipeds** | 129 | 5.968 | 5.745 – 6.191 | 0.279 | 0.145 – 0.413 | 1.093 | 0.879 – 1.307 | 0.934 | × |
| **Family** | | | | | | | | | |
| Canidae | 17 | 5.469 | 5.229 – 5.709 | 0.396 | 0.145 – 0.646 | 0.986 | 0.508 – 1.464 | 0.934 | × |
| Mustelidae | 32 | 4.631 | 4.420 – 4.842 | 0.128 | -0.014 – 0.270 | 1.417 | 0.767 – 2.067 | 0.932 | × |
| Procyonidae | 7 | 4.738 | 4.411 – 5.064 | 0.308 | -0.103 – 0.719 | 0.737 | -0.504 – 1.977 | 0.919 | × |
| Ursidae | 7 | 5.922 | 5.741 – 6.104 | 0.217 | 0.002 – 0.431 | 1.497 | 0.251 – 2.744 | 0.957 | × |
| Felidae | 26 | 5.785 | 5.691 – 5.880 | 0.232 | 0.151 – 0.314 | 1.136 | 0.927 – 1.346 | 0.987 | × |
| Herpestidae | 11 | 4.590 | 4.376 – 4.804 | 0.484 | 0.235 – 0.734 | 0.555 | 0.191 – 0.919 | 0.973 | ✓ ($D$<1) |
| Eupleridae | 5 | 4.831 | 4.633 – 5.028 | 0.317 | 0.063 – 0.570 | 1.165 | 0.309 – 2.021 | 0.996 | × |
| Viverridae | 14 | 4.922 | 4.769 – 5.076 | 0.376 | 0.195 – 0.557 | 0.565 | 0.239 – 0.891 | 0.960 | ✓ ($D$<1) |
| **Locomotor type** | | | | | | | | | |
| arboreal | 7 | 4.922 | 4.556 – 5.288 | 0.293 | -0.153 – 0.738 | 0.883 | -0.785 – 2.550 | 0.884 | × |
| semiarboreal | 10 | 5.072 | 4.939 – 5.206 | 0.410 | 0.255 – 0.565 | 0.710 | 0.450 – 0.970 | 0.987 | ✓ ($D$<1) |
| scansorial | 45 | 5.829 | 5.725 – 5.933 | 0.199 | 0.125 – 0.273 | 1.245 | 1.048 – 1.442 | 0.984 | ✓ ($D$>1) |
| terrestrial | 48 | 6.069 | 5.764 – 6.375 | 0.292 | 0.130 – 0.455 | 1.114 | 0.875 – 1.352 | 0.965 | × |
| semifossorial | 7 | 4.573 | 4.324 – 4.823 | 0.211 | -0.164 – 0.587 | 1.361 | -0.289 – 3.011 | 0.960 | × |
| semiaquatic | 11 | – | – | – | – | – | – | – | n.c. |
| aquatic | 8 | – | – | – | – | – | – | – | n.c. |

| **SR15 – N** | **n** | ***ln A*** | **95% $CI_{ln\ A}$** | ***C*** | **95% $CI_C$** | ***D*** | **95% $CI_D$** | **R** | **$D \neq 1$** |
|---|---|---|---|---|---|---|---|---|---|
| **whole sample** | 136 | 3.735 | 3.583 – 3.888 | 0.153 | 0.083 – 0.222 | 1.328 | 1.119 – 1.538 | 0.950 | ✓ ($D$>1) |
| **fissipeds** | 129 | 4.146 | 3.991 – 4.301 | 0.429 | 0.321 – 0.537 | 0.925 | 0.817 – 1.032 | 0.977 | × |
| **Family** | | | | | | | | | |
| Canidae | 17 | 3.258 | 3.036 – 3.480 | 0.362 | 0.133 – 0.590 | 1.046 | 0.562 – 1.530 | 0.938 | × |
| Mustelidae | 32 | 3.035 | 2.847 – 3.223 | 0.347 | 0.179 – 0.514 | 0.980 | 0.719 – 1.242 | 0.964 | × |
| Procyonidae | 7 | 2.706 | 2.389 – 3.022 | 0.356 | -0.085 – 0.798 | 0.967 | -0.410 – 2.343 | 0.942 | × |
| Ursidae | 7 | 3.992 | 3.702 – 4.281 | 0.166 | -0.181 – 0.514 | 1.185 | -1.352 – 3.722 | 0.809 | × |
| Felidae | 26 | 3.764 | 3.652 – 3.875 | 0.398 | 0.290 – 0.507 | 0.901 | 0.747 – 1.054 | 0.989 | × |
| Herpestidae | 11 | 2.607 | 2.445 – 2.770 | 0.543 | 0.355 – 0.731 | 0.525 | 0.291 – 0.759 | 0.988 | ✓ ($D$<1) |
| Eupleridae | 5 | 2.730 | 2.569 – 2.891 | 0.410 | 0.190 – 0.631 | 1.004 | 0.429 – 1.578 | 0.998 | × |
| Viverridae | 14 | 3.105 | 2.574 – 3.637 | 0.737 | 0.143 – 1.332 | 0.348 | -0.029 – 0.725 | 0.936 | ✓ ($D$=0) |
| **Locomotor type** | | | | | | | | | |
| arboreal | 7 | 3.033 | 2.933 – 3.132 | 0.539 | 0.407 – 0.672 | 0.664 | 0.410 – 0.919 | 0.996 | ✓ ($D$<1) |
| semiarboreal | 10 | 2.833 | 2.625 – 3.041 | 0.324 | 0.092 – 0.556 | 0.938 | 0.404 – 1.472 | 0.968 | × |
| scansorial | 45 | 3.968 | 3.824 – 4.113 | 0.363 | 0.242 – 0.484 | 1.006 | 0.837 – 1.174 | 0.984 | × |
| terrestrial | 48 | 4.140 | 3.875 – 4.405 | 0.421 | 0.253 – 0.590 | 0.940 | 0.776 – 1.105 | 0.974 | × |
| semifossorial | 7 | 2.770 | 2.539 – 3.001 | 0.236 | -0.114 – 0.585 | 1.352 | -0.025 – 2.729 | 0.972 | × |
| semiaquatic | 11 | 2.837 | 2.641 – 3.032 | 0.188 | 0.011 – 0.365 | 1.347 | 0.675 – 2.020 | 0.965 | × |
| aquatic | 8 | – | – | – | – | – | – | – | n.c. |

| SR16 – $d_{sf}$ | n | *ln A* | 95% $CI_{ln A}$ | *C* | 95% $CI_C$ | *D* | 95% $CI_D$ | R | *D* ≠ 1 |
|---|---|---|---|---|---|---|---|---|---|
| **whole sample** | 136 | 3.398 | 3.250 – 3.546 | 0.220 | 0.137 – 0.302 | 1.152 | 0.986 – 1.318 | 0.962 | × |
| **fissipeds** | 129 | 3.520 | 3.368 – 3.672 | 0.353 | 0.251 – 0.455 | 0.975 | 0.850 – 1.100 | 0.972 | × |
| **Family** | | | | | | | | | |
| Canidae | 17 | 2.785 | 2.587 – 2.983 | 0.371 | 0.167 – 0.575 | 1.044 | 0.623 – 1.465 | 0.951 | × |
| Mustelidae | 32 | 2.623 | 2.436 – 2.809 | 0.461 | 0.279 – 0.643 | 0.775 | 0.576 – 0.974 | 0.968 | ✓ (*D*<1) |
| Procyonidae | 7 | 2.263 | 2.068 – 2.459 | 0.311 | 0.037 – 0.585 | 0.979 | -0.007 – 1.965 | 0.970 | × |
| Ursidae | 7 | 3.431 | 3.159 – 3.703 | 0.262 | -0.064 – 0.588 | 1.172 | -0.335 – 2.680 | 0.917 | × |
| Felidae | 26 | 3.260 | 3.184 – 3.336 | 0.274 | 0.207 – 0.340 | 1.110 | 0.967 – 1.254 | 0.993 | × |
| Herpestidae | 11 | 2.050 | 1.897 – 2.204 | 0.336 | 0.145 – 0.527 | 0.891 | 0.359 – 1.423 | 0.963 | × |
| Eupleridae | 5 | 2.470 | 2.109 – 2.831 | 0.415 | -0.064 – 0.894 | 1.085 | -0.150 – 2.320 | 0.991 | × |
| Viverridae | 14 | 2.513 | 2.274 – 2.752 | 0.495 | 0.213 – 0.777 | 0.567 | 0.179 – 0.954 | 0.895 | ✓ (*D*<1) |
| **Locomotor type** | | | | | | | | | |
| arboreal | 7 | 2.521 | 2.272 – 2.770 | 0.419 | 0.094 – 0.743 | 0.709 | -0.111 – 1.530 | 0.966 | × |
| semiarboreal | 10 | 2.477 | 2.205 – 2.748 | 0.321 | 0.023 – 0.618 | 1.015 | 0.308 – 1.722 | 0.953 | × |
| scansorial | 45 | 3.335 | 3.249 – 3.422 | 0.252 | 0.186 – 0.318 | 1.144 | 1.007 – 1.281 | 0.991 | ✓ (*D*>1) |
| terrestrial | 48 | 3.531 | 3.264 – 3.797 | 0.382 | 0.214 – 0.551 | 0.945 | 0.763 – 1.126 | 0.969 | × |
| semifossorial | 7 | 2.160 | 1.932 – 2.389 | 0.243 | -0.154 – 0.639 | 1.103 | -0.381 – 2.586 | 0.964 | × |
| semiaquatic | 11 | 2.372 | 2.119 – 2.624 | 0.212 | -0.027 – 0.452 | 1.188 | 0.400 – 1.976 | 0.939 | × |
| aquatic | 8 | 3.707 | 2.063 – 5.351 | 0.830 | -0.975 – 2.635 | 0.336 | -0.687 – 1.359 | 0.820 | × |

| SR17 – $d_{tf}$ | n | *ln A* | 95% $CI_{ln A}$ | *C* | 95% $CI_C$ | *D* | 95% $CI_D$ | R | *D* ≠ 1 |
|---|---|---|---|---|---|---|---|---|---|
| **whole sample** | 136 | 3.942 | 3.756 – 4.127 | 0.459 | 0.326 – 0.592 | 0.880 | 0.761 – 0.999 | 0.973 | ✓ (*D*<1) |
| **fissipeds** | 129 | 3.665 | 3.491 – 3.838 | 0.387 | 0.268 – 0.506 | 0.947 | 0.815 – 1.079 | 0.967 | × |
| **Family** | | | | | | | | | |
| Canidae | 17 | 2.794 | 2.602 – 2.986 | 0.361 | 0.164 – 0.558 | 1.055 | 0.635 – 1.474 | 0.952 | × |
| Mustelidae | 32 | 2.787 | 2.581 – 2.993 | 0.500 | 0.301 – 0.699 | 0.797 | 0.595 – 1.000 | 0.969 | × |
| Procyonidae | 7 | 2.361 | 2.141 – 2.581 | 0.334 | 0.052 – 0.616 | 0.779 | -0.038 – 1.595 | 0.966 | × |
| Ursidae | 7 | 3.483 | 3.199 – 3.767 | 0.112 | -0.216 – 0.441 | 1.698 | -2.038 – 5.433 | 0.773 | × |
| Felidae | 26 | 3.350 | 3.210 – 3.491 | 0.299 | 0.173 – 0.425 | 1.066 | 0.819 – 1.313 | 0.979 | × |
| Herpestidae | 11 | 2.129 | 1.915 – 2.343 | 0.352 | 0.092 – 0.611 | 0.703 | 0.090 – 1.316 | 0.941 | × |
| Eupleridae | 5 | 2.392 | 2.260 – 2.523 | 0.336 | 0.153 – 0.519 | 0.974 | 0.391 – 1.557 | 0.998 | × |
| Viverridae | 14 | 2.637 | 2.360 – 2.913 | 0.551 | 0.230 – 0.872 | 0.449 | 0.111 – 0.788 | 0.952 | ✓ (*D*<1) |
| **Locomotor type** | | | | | | | | | |
| arboreal | 7 | 2.691 | 2.438 – 2.945 | 0.565 | 0.236 – 0.894 | 0.418 | -0.018 – 0.854 | 0.991 | ✓ (*D*=0) |
| semiarboreal | 10 | 2.564 | 2.317 – 2.811 | 0.407 | 0.124 – 0.690 | 0.782 | 0.289 – 1.275 | 0.962 | × |
| scansorial | 45 | 3.549 | 3.401 – 3.696 | 0.345 | 0.221 – 0.470 | 0.995 | 0.814 – 1.177 | 0.982 | × |
| terrestrial | 48 | 3.552 | 3.269 – 3.835 | 0.371 | 0.195 – 0.548 | 0.957 | 0.761 – 1.154 | 0.965 | × |
| semifossorial | 7 | 2.247 | 2.005 – 2.488 | 0.234 | -0.168 – 0.637 | 1.191 | -0.386 – 2.768 | 0.961 | × |
| semiaquatic | 11 | 2.603 | 2.289 – 2.916 | 0.296 | -0.015 – 0.607 | 1.029 | 0.319 – 1.739 | 0.932 | × |
| aquatic | 8 | 4.109 | 3.417 – 4.801 | 0.675 | -0.092 – 1.442 | 0.644 | -0.128 – 1.415 | 0.901 | × |

| **SR18 – FR** | **n** | ***ln A*** | **95% CI$_{ln A}$** | ***C*** | **95% CI$_C$** | ***D*** | **95% CI$_D$** | **R** | ***D* ≠ 1** |
|---|---|---|---|---|---|---|---|---|---|
| **whole sample** | 136 | – | – | – | – | – | – | – | n.c. |
| **fissipeds** | 129 | -2.542 | -2.582 – -2.503 | $-7.87 \cdot 10^{-7}$ | $-1.30 \cdot 10^{-5} – 1.15 \cdot 10^{-5}$ | 6.052 | -1.755 – 13.858 | 0.197 | × |
| **Family** | | | | | | | | | |
| Canidae | 17 | – | – | – | – | – | – | – | n.c. |
| Mustelidae | 32 | – | – | – | – | – | – | – | n.c. |
| Procyonidae | 7 | – | – | – | – | – | – | – | n.c. |
| Ursidae | 7 | – | – | – | – | – | – | – | n.c. |
| Felidae | 26 | -2.529 | -2.639 – -2.418 | 0.038 | -0.065 – 0.140 | 1.000 | -0.573 – 2.573 | 0.587 | × |
| Herpestidae | 11 | – | – | – | – | – | – | – | n.c. |
| Eupleridae | *5* | *-2.343* | *-2.841 – -1.845* | *0.124* | *-0.650 – 0.897* | *0.604* | *-4.965 – 6.173* | *0.840* | n.s. |
| Viverridae | 14 | -2.405 | -2.543 – -2.266 | 0.122 | -0.041 – 0.286 | 0.568 | -0.341 – 1.476 | 0.777 | × |
| **Locomotor type** | | | | | | | | | |
| arboreal | 7 | – | – | – | – | – | – | – | n.c. |
| semiarboreal | 10 | – | – | – | – | – | – | – | n.c. |
| scansorial | *45* | *-2.502* | *-2.718 – -2.285* | *0.049* | *-0.173 – 0.271* | *0.504* | *-1.226 – 2.234* | *0.288* | n.s. |
| terrestrial | 48 | -2.654 | -2.787 – -2.521 | -0.003 | -0.021 – 0.015 | 2.236 | -0.296 – 4.768 | 0.558 | × |
| semifossorial | 7 | – | – | – | – | – | – | – | n.c. |
| semiaquatic | 11 | -2.133 | -2.366 – -1.899 | 0.137 | -0.112 – 0.385 | 0.718 | -0.395 – 1.831 | 0.762 | × |
| aquatic | 8 | – | – | – | – | – | – | – | n.c. |

| **SR19 – $L_t$** | **n** | ***ln A*** | **95% CI$_{ln A}$** | ***C*** | **95% CI$_C$** | ***D*** | **95% CI$_D$** | **R** | ***D* ≠ 1** |
|---|---|---|---|---|---|---|---|---|---|
| **whole sample** | 136 | 5.628 | 5.474 – 5.782 | 0.097 | 0.039 – 0.156 | 1.486 | 1.205 – 1.766 | 0.953 | ✓ (*D*>1) |
| **fissipeds** | 129 | 5.643 | 5.456 – 5.830 | 0.146 | 0.058 – 0.234 | 1.332 | 1.051 – 1.613 | 0.915 | ✓ (*D*>1) |
| **Family** | | | | | | | | | |
| Canidae | 17 | 5.504 | 5.163 – 5.845 | 0.407 | 0.043 – 0.770 | 0.881 | 0.229 – 1.533 | 0.875 | × |
| Mustelidae | 32 | 4.783 | 4.524 – 5.041 | 0.222 | 0.004 – 0.439 | 1.082 | 0.539 – 1.626 | 0.885 | × |
| Procyonidae | 7 | 4.818 | 4.307 – 5.328 | 0.394 | -0.171 – 0.960 | 0.452 | -0.451 – 1.355 | 0.917 | × |
| Ursidae | 7 | 5.637 | 5.497 – 5.777 | 0.206 | 0.042 – 0.371 | 1.558 | 0.551 – 2.566 | 0.973 | × |
| Felidae | 26 | 5.622 | 5.494 – 5.749 | 0.162 | 0.060 – 0.265 | 1.252 | 0.867 – 1.636 | 0.964 | × |
| Herpestidae | 11 | 4.687 | 4.283 – 5.092 | 0.547 | 0.091 – 1.003 | 0.454 | -0.045 – 0.953 | 0.942 | ✓ (*D*=0) |
| Eupleridae | 5 | 4.779 | 4.358 – 5.199 | 0.200 | -0.280 – 0.679 | 1.457 | -1.103 – 4.017 | 0.972 | × |
| Viverridae | 14 | 4.847 | 4.715 – 4.979 | 0.295 | 0.139 – 0.451 | 0.611 | 0.237 – 0.986 | 0.951 | ✓ (*D*<1) |
| **Locomotor type** | | | | | | | | | |
| arboreal | 7 | 4.851 | 4.362 – 5.340 | 0.257 | -0.346 – 0.860 | 0.846 | -1.714 – 3.406 | 0.775 | × |
| semiarboreal | 10 | 4.971 | 4.778 – 5.164 | 0.343 | 0.119 – 0.567 | 0.670 | 0.229 – 1.111 | 0.962 | × |
| scansorial | 45 | 5.552 | 5.434 – 5.669 | 0.105 | 0.037 – 0.173 | 1.478 | 1.124 – 1.831 | 0.960 | ✓ (*D*>1) |
| terrestrial | 48 | 5.794 | 5.440 – 6.147 | 0.179 | 0.023 – 0.335 | 1.290 | 0.905 – 1.674 | 0.939 | × |
| semifossorial | 7 | – | – | – | – | – | – | – | n.c. |
| semiaquatic | 11 | 4.657 | 4.377 – 4.936 | 0.101 | -0.130 – 0.332 | 1.586 | -0.103 – 3.275 | 0.872 | × |
| aquatic | 8 | 5.699 | 5.527 – 5.870 | 0.259 | 0.078 – 0.440 | 1.066 | 0.533 – 1.599 | 0.976 | × |

| SR20 – $d_{st}$ | n | *ln A* | 95% $CI_{ln A}$ | *C* | 95% $CI_C$ | *D* | 95% $CI_D$ | R | *D* ≠ 1 |
|---|---|---|---|---|---|---|---|---|---|
| **whole sample** | 136 | 3.385 | 3.237 – 3.533 | 0.170 | 0.098 – 0.242 | 1.278 | 1.086 – 1.470 | 0.956 | ✓ (*D*>1) |
| **fissipeds** | 129 | 3.676 | 3.516 – 3.836 | 0.392 | 0.283 – 0.501 | 0.950 | 0.830 – 1.070 | 0.973 | × |
| **Family** | | | | | | | | | |
| Canidae | 17 | 2.876 | 2.704 – 3.048 | 0.428 | 0.246 – 0.609 | 0.936 | 0.621 – 1.251 | 0.968 | × |
| Mustelidae | 32 | 2.708 | 2.494 – 2.922 | 0.436 | 0.235 – 0.638 | 0.852 | 0.611 – 1.092 | 0.961 | × |
| Procyonidae | 7 | 2.248 | 1.842 – 2.653 | 0.296 | -0.213 – 0.804 | 0.731 | -0.859 – 2.321 | 0.875 | × |
| Ursidae | 7 | 3.505 | 3.196 – 3.814 | 0.179 | -0.167 – 0.525 | 1.968 | -0.525 – 4.462 | 0.901 | × |
| Felidae | 26 | 3.362 | 3.246 – 3.477 | 0.243 | 0.149 – 0.338 | 1.232 | 0.998 – 1.467 | 0.986 | × |
| Herpestidae | 11 | 2.139 | 1.955 – 2.322 | 0.390 | 0.173 – 0.606 | 0.592 | 0.179 – 1.005 | 0.967 | × |
| Eupleridae | 5 | 2.478 | 2.274 – 2.682 | 0.520 | 0.204 – 0.835 | 0.732 | 0.105 – 1.359 | 0.997 | × |
| Viverridae | 14 | 2.482 | 2.369 – 2.596 | 0.427 | 0.294 – 0.561 | 0.634 | 0.409 – 0.859 | 0.982 | ✓ (*D*<1) |
| **Locomotor type** | | | | | | | | | |
| arboreal | 7 | 2.508 | 2.298 – 2.717 | 0.362 | 0.108 – 0.615 | 0.897 | 0.128 – 1.665 | 0.972 | × |
| semiarboreal | 10 | 2.637 | 2.428 – 2.847 | 0.423 | 0.189 – 0.658 | 0.931 | 0.519 – 1.343 | 0.980 | × |
| scansorial | 45 | 3.485 | 3.355 – 3.615 | 0.292 | 0.188 – 0.395 | 1.090 | 0.908 – 1.273 | 0.984 | × |
| terrestrial | 48 | 3.665 | 3.398 – 3.931 | 0.426 | 0.252 – 0.600 | 0.909 | 0.742 – 1.076 | 0.971 | × |
| semifossorial | 7 | 2.308 | 2.055 – 2.561 | 0.294 | -0.148 – 0.737 | 1.084 | -0.278 – 2.446 | 0.969 | × |
| semiaquatic | 11 | 2.569 | 2.224 – 2.913 | 0.318 | -0.026 – 0.662 | 0.999 | 0.273 – 1.726 | 0.925 | × |
| aquatic | 8 | – | – | – | – | – | – | – | n.c. |

| SR21 – $d_{tt}$ | n | *ln A* | 95% $CI_{ln A}$ | *C* | 95% $CI_C$ | *D* | 95% $CI_D$ | R | *D* ≠ 1 |
|---|---|---|---|---|---|---|---|---|---|
| **whole sample** | 136 | 3.484 | 3.312 – 3.656 | 0.287 | 0.184 – 0.390 | 1.078 | 0.922 – 1.235 | 0.964 | × |
| **fissipeds** | 129 | 3.388 | 3.208 – 3.568 | 0.316 | 0.204 – 0.429 | 1.048 | 0.891 – 1.206 | 0.961 | × |
| **Family** | | | | | | | | | |
| Canidae | 17 | 2.830 | 2.603 – 3.057 | 0.442 | 0.202 – 0.682 | 0.914 | 0.512 – 1.315 | 0.921 | × |
| Mustelidae | 32 | 2.393 | 2.188 – 2.599 | 0.395 | 0.203 – 0.587 | 0.878 | 0.623 – 1.133 | 0.959 | × |
| Procyonidae | 7 | 2.042 | 1.767 – 2.318 | 0.359 | -0.014 – 0.733 | 0.904 | -0.204 – 2.012 | 0.955 | × |
| Ursidae | 7 | 3.205 | 2.905 – 3.505 | 0.148 | -0.196 – 0.492 | 1.785 | -1.195 – 4.766 | 0.846 | × |
| Felidae | 26 | 3.233 | 3.107 – 3.360 | 0.296 | 0.184 – 0.408 | 1.087 | 0.864 – 1.311 | 0.983 | × |
| Herpestidae | 11 | 2.101 | 1.591 – 2.610 | 0.556 | 0.001 – 1.111 | 0.363 | -0.120 – 0.847 | 0.938 | ✓ (*D*=0) |
| Eupleridae | 5 | 2.110 | 1.507 – 2.713 | 0.250 | -0.459 – 0.958 | 1.383 | -1.644 – 4.410 | 0.959 | × |
| Viverridae | 14 | 2.258 | 2.169 – 2.347 | 0.376 | 0.273 – 0.479 | 0.810 | 0.594 – 1.026 | 0.986 | × |
| **Locomotor type** | | | | | | | | | |
| arboreal | 7 | 2.248 | 1.953 – 2.544 | 0.313 | -0.019 – 0.645 | 1.111 | -0.055 – 2.277 | 0.946 | × |
| semiarboreal | 10 | 2.252 | 2.040 – 2.465 | 0.327 | 0.090 – 0.565 | 0.932 | 0.392 – 1.473 | 0.967 | × |
| scansorial | 45 | 3.243 | 3.122 – 3.365 | 0.240 | 0.150 – 0.329 | 1.191 | 0.995 – 1.388 | 0.983 | × |
| terrestrial | 48 | 3.336 | 3.008 – 3.664 | 0.277 | 0.101 – 0.452 | 1.111 | 0.840 – 1.382 | 0.955 | × |
| semifossorial | 7 | 1.968 | 1.512 – 2.424 | 0.401 | -0.372 – 1.175 | 0.740 | -0.797 – 2.276 | 0.948 | × |
| semiaquatic | 11 | 2.369 | 2.092 – 2.646 | 0.372 | 0.085 – 0.659 | 0.838 | 0.343 – 1.333 | 0.949 | × |
| aquatic | 8 | 3.618 | 3.018 – 4.217 | 0.516 | -0.132 – 1.164 | 0.863 | -0.054 – 1.779 | 0.903 | × |

| SR22 – TR | n | *ln A* | 95% CI$_{ln A}$ | *C* | 95% CI$_C$ | *D* | 95% CI$_D$ | R | *D* ≠ 1 |
|---|---|---|---|---|---|---|---|---|---|
| **whole sample** | 136 | -2.305 | -2.557 – -2.053 | 0.048 | -0.137 – 0.233 | 0.851 | -0.703 – 2.405 | 0.298 | × |
| **fissipeds** | 129 | -1.865 | -2.584 – -1.147 | 0.414 | -0.308 – 1.137 | 0.232 | -0.131 – 0.696 | 0.451 | ✓ (*D*=0) |
| **Family** | | | | | | | | | |
| Canidae | 17 | – | – | – | – | – | – | – | n.c. |
| Mustelidae | 32 | -2.104 | -2.417 – -1.790 | 0.184 | -0.142 – 0.510 | 0.563 | -0.215 – 1.340 | 0.628 | × |
| Procyonidae | 7 | – | – | – | – | – | – | – | n.c. |
| Ursidae | 7 | – | – | – | – | – | – | – | n.c. |
| Felidae | 26 | -2.255 | -2.438 – -2.073 | 0.084 | -0.070 – 0.239 | 1.167 | 0.073 – 2.261 | 0.768 | × |
| Herpestidae | 11 | – | – | – | – | – | – | – | n.c. |
| Eupleridae | 5 | – | – | – | – | – | – | – | n.c. |
| Viverridae | 14 | -2.366 | -2.502 – -2.230 | 0.130 | -0.030 – 0.290 | 0.701 | -0.225 – 1.628 | 0.794 | × |
| **Locomotor type** | | | | | | | | | |
| arboreal | 7 | – | – | – | – | – | – | – | n.c. |
| semiarboreal | 10 | -2.302 | -2.496 – -2.108 | 0.125 | -0.075 – 0.325 | 1.246 | -0.029 – 2.522 | 0.907 | × |
| scansorial | 45 | -2.071 | -2.333 – -1.810 | 0.198 | -0.061 – 0.457 | 0.645 | 0.082 – 1.207 | 0.756 | × |
| terrestrial | 48 | – | – | – | – | – | – | – | n.c. |
| semifossorial | 7 | -1.894 | -2.766 – -1.023 | 0.438 | -0.485 – 1.360 | 0.195 | -0.307 – 0.698 | 0.976 | ✓ (*D*=0) |
| semiaquatic | 11 | -2.155 | -2.482 – -1.828 | 0.143 | -0.203 – 0.490 | 0.733 | -0.759 – 2.225 | 0.665 | × |
| aquatic | 8 | – | – | – | – | – | – | – | n.c. |

| SR23 – L$_c$ | n | *ln A* | 95% CI$_{ln A}$ | *C* | 95% CI$_C$ | *D* | 95% CI$_D$ | R | *D* ≠ 1 |
|---|---|---|---|---|---|---|---|---|---|
| **whole sample** | 85 | 4.518 | 4.363 – 4.672 | 0.149 | 0.082 – 0.215 | 1.359 | 1.153 – 1.565 | 0.962 | ✓ (*D*>1) |
| **fissipeds** | 84 | 4.531 | 4.377 – 4.684 | 0.220 | 0.136 – 0.305 | 1.204 | 1.024 – 1.383 | 0.964 | ✓ (*D*>1) |
| **Family** | | | | | | | | | |
| Canidae | 19 | 4.089 | 3.859 – 4.319 | 0.434 | 0.193 – 0.675 | 0.955 | 0.539 – 1.371 | 0.943 | × |
| Mustelidae | 15 | 3.475 | 3.262 – 3.688 | 0.436 | 0.196 – 0.676 | 0.812 | 0.477 – 1.148 | 0.961 | × |
| Procyonidae | | | | | | | | | |
| Ursidae | 5 | 4.530 | 3.650 – 5.409 | 0.424 | -0.599 – 1.447 | 0.722 | -3.029 – 4.473 | 0.887 | × |
| Felidae | 21 | 4.533 | 4.415 – 4.651 | 0.334 | 0.216 – 0.453 | 0.971 | 0.748 – 1.193 | 0.984 | × |
| Herpestidae | | | | | | | | | |
| Eupleridae | | | | | | | | | |
| Viverridae | 9 | 3.529 | 3.324 – 3.733 | 0.339 | 0.092 – 0.586 | 0.775 | 0.190 – 1.361 | 0.952 | × |
| **Locomotor type** | | | | | | | | | |
| arboreal | 6 | 3.634 | 3.298 – 3.969 | 0.452 | -0.068 – 0.972 | 0.608 | -0.840 – 2.057 | 0.973 | × |
| semiarboreal | 5 | 3.379 | 3.016 – 3.742 | 0.255 | -0.234 – 0.744 | 1.164 | -0.745 – 3.073 | 0.975 | × |
| scansorial | 35 | 4.389 | 4.253 – 4.525 | 0.171 | 0.080 – 0.261 | 1.340 | 1.045 – 1.635 | 0.970 | ✓ (*D*>1) |
| terrestrial | 30 | 4.625 | 4.312 – 4.938 | 0.269 | 0.107 – 0.430 | 1.134 | 0.872 – 1.395 | 0.963 | × |
| semifossorial | | | | | | | | | |
| semiaquatic | 6 | – | – | – | – | – | – | – | n.c. |
| aquatic | | | | | | | | | |

| SR24 – r | n | *ln A* | 95% CI$_{ln A}$ | *C* | 95% CI$_C$ | *D* | 95% CI$_D$ | R | $D \neq 1$ |
|---|---|---|---|---|---|---|---|---|---|
| **whole sample** | 85 | 4.341 | 4.188 – 4.493 | 0.184 | 0.112 – 0.257 | 1.274 | 1.094 – 1.453 | 0.969 | ✓ (*D*>1) |
| **fissipeds** | 84 | 4.340 | 4.190 – 4.491 | 0.261 | 0.172 – 0.349 | 1.137 | 0.979 – 1.295 | 0.970 | × |
| **Family** | | | | | | | | | |
| Canidae | 19 | 3.874 | 3.631 – 4.117 | 0.473 | 0.217 – 0.730 | 0.937 | 0.534 – 1.339 | 0.946 | × |
| Mustelidae | 15 | 3.239 | 3.035 – 3.444 | 0.445 | 0.215 – 0.675 | 0.820 | 0.505 – 1.135 | 0.966 | × |
| Procyonidae | | | | | | | | | |
| Ursidae | *5* | *4.352* | *3.212 – 5.492* | *0.450* | *-0.848 – 1.748* | *0.648* | *-3.407 – 4.702* | *0.858* | n.s. |
| Felidae | 21 | 4.268 | 4.108 – 4.429 | 0.328 | 0.172 – 0.485 | 1.043 | 0.740 – 1.347 | 0.974 | × |
| Herpestidae | | | | | | | | | |
| Eupleridae | | | | | | | | | |
| Viverridae | 9 | 3.301 | 3.108 – 3.494 | 0.400 | 0.163 – 0.637 | 0.710 | 0.244 – 1.175 | 0.967 | × |
| **Locomotor type** | | | | | | | | | |
| arboreal | 6 | – | – | – | – | – | – | – | n.c. |
| semiarboreal | 5 | 3.093 | 2.724 – 3.461 | 0.262 | -0.235 – 0.760 | 1.146 | -0.739 – 3.030 | 0.975 | × |
| scansorial | 35 | 4.190 | 4.049 – 4.330 | 0.218 | 0.115 – 0.320 | 1.232 | 0.974 – 1.490 | 0.975 | × |
| terrestrial | 30 | 4.438 | 4.131 – 4.746 | 0.302 | 0.136 – 0.467 | 1.092 | 0.854 – 1.329 | 0.966 | × |
| semifossorial | | | | | | | | | |
| semiaquatic | 6 | – | – | – | – | – | – | – | n.c. |
| aquatic | | | | | | | | | |

| SR25 – d$_{sc}$ | n | *ln A* | 95% CI$_{ln A}$ | *C* | 95% CI$_C$ | *D* | 95% CI$_D$ | R | $D \neq 1$ |
|---|---|---|---|---|---|---|---|---|---|
| **whole sample** | 85 | 3.706 | 3.520 – 3.891 | 0.256 | 0.151 – 0.360 | 1.117 | 0.935 – 1.299 | 0.962 | × |
| **fissipeds** | 84 | 3.775 | 3.592 – 3.959 | 0.397 | 0.266 – 0.528 | 0.937 | 0.791 – 1.083 | 0.967 | × |
| **Family** | | | | | | | | | |
| Canidae | 19 | 3.142 | 2.772 – 3.511 | 0.585 | 0.186 – 0.984 | 0.784 | 0.308 – 1.259 | 0.912 | × |
| Mustelidae | 15 | 2.549 | 2.326 – 2.772 | 0.419 | 0.171 – 0.666 | 0.860 | 0.494 – 1.225 | 0.958 | × |
| Procyonidae | | | | | | | | | |
| Ursidae | 5 | 3.743 | 2.870 – 4.617 | 0.473 | -0.586 – 1.532 | 0.886 | -3.255 – 5.026 | 0.897 | × |
| Felidae | 21 | 3.572 | 3.400 – 3.743 | 0.404 | 0.229 – 0.580 | 0.918 | 0.650 – 1.187 | 0.974 | × |
| Herpestidae | | | | | | | | | |
| Eupleridae | | | | | | | | | |
| Viverridae | 9 | 2.554 | 2.408 – 2.699 | 0.363 | 0.185 – 0.540 | 0.737 | 0.350 – 1.125 | 0.977 | × |
| **Locomotor type** | | | | | | | | | |
| arboreal | 6 | – | – | – | – | – | – | – | n.c. |
| semiarboreal | 5 | 2.456 | 2.101 – 2.812 | 0.369 | -0.108 – 0.845 | 0.828 | -0.358 – 2.015 | 0.983 | × |
| scansorial | 35 | 3.595 | 3.450 – 3.741 | 0.349 | 0.222 – 0.477 | 0.971 | 0.782 – 1.160 | 0.981 | × |
| terrestrial | 30 | 3.839 | 3.482 – 4.195 | 0.435 | 0.205 – 0.666 | 0.919 | 0.695 – 1.144 | 0.954 | × |
| semifossorial | | | | | | | | | |
| semiaquatic | 6 | – | – | – | – | – | – | – | n.c. |
| aquatic | | | | | | | | | |

| **SR26 – $d_{tc}$** | **n** | ***ln A*** | **95% $CI_{ln A}$** | ***C*** | **95% $CI_C$** | ***D*** | **95% $CI_D$** | **R** | ***D* ≠ 1** |
|---|---|---|---|---|---|---|---|---|---|
| **whole sample** | 85 | 3.058 | 2.835 – 3.282 | 0.312 | 0.173 – 0.451 | 1.024 | 0.829 – 1.218 | 0.952 | × |
| **fissipeds** | 84 | 2.905 | 2.696 – 3.114 | 0.317 | 0.179 – 0.456 | 1.018 | 0.820 – 1.215 | 0.948 | × |
| **Family** | | | | | | | | | |
| Canidae | 19 | 2.252 | 1.988 – 2.516 | 0.361 | 0.091 – 0.632 | 1.052 | 0.475 – 1.629 | 0.909 | × |
| Mustelidae | 15 | 1.783 | 1.519 – 2.047 | 0.419 | 0.123 – 0.716 | 0.819 | 0.386 – 1.251 | 0.939 | × |
| Procyonidae | | | | | | | | | |
| Ursidae | 5 | – | – | – | – | – | – | – | n.c. |
| Felidae | 21 | 2.825 | 2.623 – 3.026 | 0.374 | 0.169 – 0.579 | 0.937 | 0.597 – 1.277 | 0.961 | × |
| Herpestidae | | | | | | | | | |
| Eupleridae | | | | | | | | | |
| Viverridae | 9 | 1.940 | 1.833 – 2.047 | 0.504 | 0.371 – 0.637 | 0.513 | 0.333 – 0.692 | 0.994 | ✓ (*D*<1) |
| **Locomotor type** | | | | | | | | | |
| arboreal | 6 | – | – | – | – | – | – | – | n.c. |
| semiarboreal | 5 | – | – | – | – | – | – | – | n.c. |
| scansorial | 35 | 2.695 | 2.465 – 2.925 | 0.235 | 0.056 – 0.414 | 1.146 | 0.735 – 1.557 | 0.934 | × |
| terrestrial | 30 | 3.051 | 2.701 – 3.400 | 0.396 | 0.177 – 0.615 | 0.949 | 0.714 – 1.184 | 0.954 | × |
| semifossorial | | | | | | | | | |
| semiaquatic | 6 | – | – | – | – | – | – | – | n.c. |
| aquatic | | | | | | | | | |
